# Emergent orientation selectivity from random networks in mouse visual cortex

**DOI:** 10.1101/285197

**Authors:** J.J. Pattadkal, G. Mato, C. van Vreeswijk, N. J. Priebe, D. Hansel

## Abstract

We study the connectivity principles underlying the emergence of orientation selectivity in primary visual cortex (V1) of mammals lacking an orientation map. We present a computational model in which random connectivity gives rise to orientation selectivity that matches experimental observations. It predicts that mouse V1 neurons should exhibit intricate receptive fields in the two-dimensional frequency domain, causing shift in orientation preferences with spatial frequency. We find evidence for these features in mouse V1 using calcium imaging and intracellular whole cell recordings.

## Introduction

Since its initial description by Hubel and Wiesel (Hubel and Wiesel, 1962), orientation selectivity has served as a platform for studying neocortical computations (Priebe and Ferster, 2012). V1 neurons in primates and carnivores are characterized not only by their preference for the orientation of bars or edges, but also that the preference for a bar or edge of a specific orientation is invariant to the spatial structure of the object displayed. For example, a V1 neuron which responds best to a vertical orientation should maintain that orientation preference despite changes in the width or movement of a presented bar (De Valois et al., 1982; Jones et al., 1987; Webster and De Valois, 1985).

Orientation selectivity emerges in V1 of primates and carnivores where a functional organization for this selectivity is also observed: neurons are organized in a columnar fashion with shared orientation preference across cortical layers and smooth changes in selectivity along the V1 surface (Hubel and Wiesel, 1977). This functional architecture is the product of the spatial arrangement of ON and OFF thalamocortical inputs that innervate V1 (Kremkow et al., 2016; Lee et al., 2016a) and of the vertical bias of intracortical connectivity (Song et al., 2005). These spatially offset ON and OFF afferents converge on individual V1 neurons to generate receptive fields that are orientation-tuned (Alonso et al., 2001) and well-described by Gabor functions (Jones and Palmer, 1987) (Fig. 1A).

**Figure 1:**
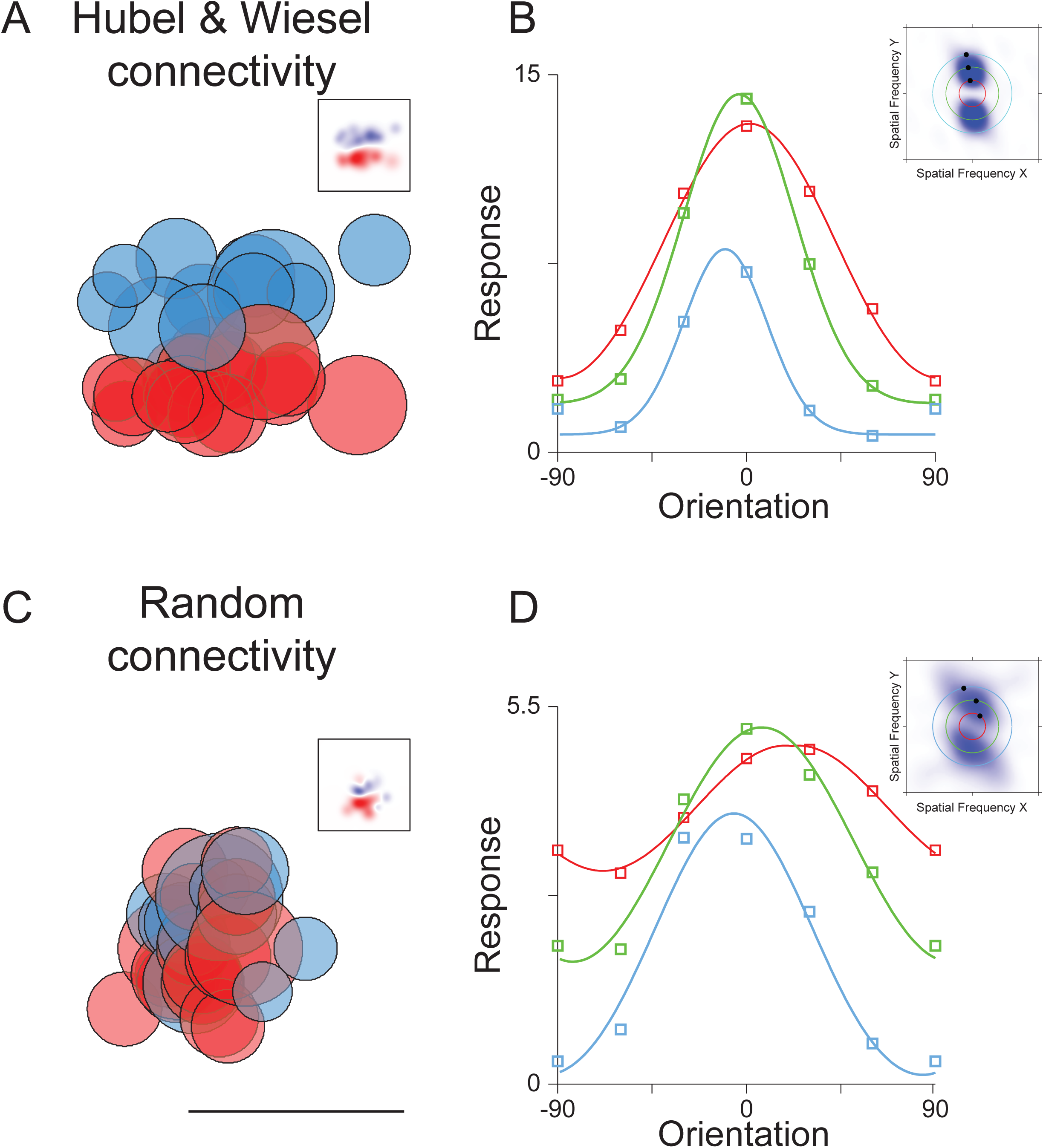
Receptive fields, random connectivity, spatial frequency tuning and orientation tuning. **A.** Hubel and Wiesel connectivity in which ON (red) and OFF (blue) thalamocortical afferents, with spatial receptive fields indicated by each circle, converge onto a neuron in primary visual cortex. The summation of these afferent receptive fields generates a Gabor like receptive field in visual cortex (inset). **B.** Orientation preference does not change with spatial frequency for such receptive fields. Tuning curves of the temporal modulation of the response for tow (red) medium (green) and high spatial frequencies are plotted. In frequency space these receptive fields maintain a peak response at a consistent angle that points toward the origin at the midpoint of the graph (inset). **C.** Random connectivity from the LGN in which ON and OFF thalamocortical neurons with similar spatial receptive fields converge on cortical neurons also generates orientation selectivity in the temporal modulation of the response. The summation of LGN neuron receptive fields shows oriented profiles (inset). Scale bar indicates 35 degrees. **D.** Orientation preference shifts for random connectivity as spatial frequency changes. Orientation tuning curves are plotted as in B. In frequency space these receptive fields tilt in a manner that does not project back to the origin.

Such a functional architecture for orientation selectivity, however, is not common to all mammals: V1 of rodents and lagomorphs lack it but their neurons are still orientation selective (Drager, 1975; Girman et al., 1999; Metin et al., 1988; Murphy and Berman, 1979). This raises the question of what connectivity rules guide afferent and intracortical circuitry to generate orientation selectivity in mammals that lack a functional architecture for orientation selectivity (Ohki and Reid, 2007).

We recently showed in a model of rodent V1 that layer 2/3 can inherit orientation selectivity from orientation selective neurons in layer 4 even if recurrent as well as feedforward (L4 to L2/3) connectivity is random (Hansel and van Vreeswijk, 2012). In this model the L2/3 network operates in a ‘balanced’ regime (van Vreeswijk and Sompolinsky, 1996, 1998), in which excitatory and inhibitory inputs, are both strong, and roughly cancel each other (Hansel and van Vreeswijk, 2012; Pehlevan and Sompolinsky, 2014).

In this report we address the question of whether orientation selectivity can emerge in rodent V1 from random connectivity. We present a strongly recurrent model of the rodent V1 network in which neurons receive inputs from randomly chosen non-selective LGN cells. It does not necessitate sparse connectivity to generate selectivity, as is required in previous random network based models of orientation tuning (Ringach, 2004; Soodak, 1987; von der Malsburg, 1973). Remarkably, orientation selectivity emerges in this network despite the lack of a Gabor like structure of the thalamocortical input with well segregated ON and OFF subfields. Furthermore, orientation selectivity in this network is robust to changes in the number of inputs. A key prediction of this model is that the orientation selectivity of V1 neurons may vary with the spatial content of the presented stimulus (Miller, 2016). It thus predicts that in mouse V1 neuron receptive fields in the frequency domain are intricate, containing dependencies between orientation and spatial frequency, in stark contrast to observations made in primates and carnivores, and predictions of Gabor receptive fields (De Valois et al., 1982; Jones et al., 1987; Webster and De Valois, 1985). To test these predictions we quantified in mouse V1 the degree to which orientation preference is linked to the stimulus spatial frequency using a combination of electrophysiological and imaging measurements. In agreement with our model we found that orientation preference depends strongly on spatial frequency.

## Results

To contrast different circuitry that could give rise to cortical orientation selectivity we constructed two model V1 neurons that receive input from the thalamus. In one model the V1 neuron receives ON and OFF thalamic inputs that are sampled on the basis of a Gabor filter: ON and OFF inputs have spatial preferences elongated along the preferred orientation axis and are spatially segregated (Fig. 1A). The temporally modulated component (F1) of the response is largest to horizontally oriented drifting gratings regardless of the spatial frequency (Fig. 1B). We also constructed a model V1 neuron in which ON and OFF inputs with nearby spatial preferences, which are randomly intermixed (Fig. 1C). Remarkably, this random connectivity model also exhibits orientation selectivity in the F1 component of the response. It emerges from the imbalances in ON and OFF inputs onto the target neuron. Unlike the ordered receptive field neuron, however, the preferred orientation of the F1 response of the cell changes with the stimulus spatial frequency. At high spatial frequency the F1 responses of the model neuron are largest for stimuli oriented at 30 degrees while at low spatial frequency responses are largest at −10 degrees (Fig. 1D). This shift in orientation preference is a product of the random connectivity onto the neuron: the imbalances of ON and OFF thalamic inputs are different as spatial scale changes, causing shifts in orientation preference.

### Orientation selectivity emerges in a model of rodent V1 with random wiring

To study whether orientation selectivity in mouse V1 could result from random connectivity we constructed a large-scale conductance-based spiking network model of V1 (Sup. Fig. 1) in which cortical neurons receive feedforward excitation from randomly chosen thalamic relay cells as well as other cortical cells of similar retinotopic preferences (Sup. Fig. 1B; see Methods). Previously it has been shown that orientation selectivity can emerge on the basis of random inputs alone (Ringach, 2004; Soodak, 1987; von der Malsburg, 1973). Orientation selectivity arises in these models because of asymmetries in the spatial preferences of the sparse inputs that converge onto a cortical neuron. As the number of convergent inputs increases, however, the selectivity declines because the tuned temporally modulated component of the LGN input decreases relative to the time averaged untuned component. To surmount this dependence of orientation selectivity on the number of inputs we employ a network model in which excitatory and inhibitory inputs are strong but balanced (van Vreeswijk and Sompolinsky, 1996, 1998) such that the mean and variance of the net input is on the order of the distance to threshold (Sup. Fig. 2).

Networks with random connectivity operating in a balanced regime have previously been shown to maintain preferences present in the input (Hansel and van Vreeswijk, 2012). We hypothesized that orientation selectivity would emerge in our model if the spatial inhomogeneity in the aggregate thalamic input were maintained in the output by the balance of excitation and inhibition. In the balanced state the untuned time-averaged component of the input is largely suppressed by the intracortical feedback, leading to a net input in which the tuned modulation is comparable to the untuned component. Indeed orientation selectivity emerges in our model (Fig. 2A), varying between highly selective neurons (*e.g* model neuron E10371) to weakly selective (*e.g* model neuron E11763). This diversity of selectivity results in a distribution of orientation selectivity index (OSI) demonstrating that orientation selectivity emerges naturally in a random connectivity model (Fig. 2, Sup. Fig. 2B, 3,4). The emergent cortical orientation preference is matched to the preferred orientation of aggregate thalamic input (Sup. Fig. 3C,D), as observed in mouse visual cortex (Li et al., 2013). In this balanced model the emergent orientation selectivity should be insensitive to the number of inputs. To verify this we varied this number from 25 to 100 and found that the degree of orientation selectivity was maintained (Fig. 2C, D, Sup Fig. 5). The emergent selectivity is also robust to changes in network size and in synaptic strength (Sup. Fig. 5A,C).

**Figure 2:**
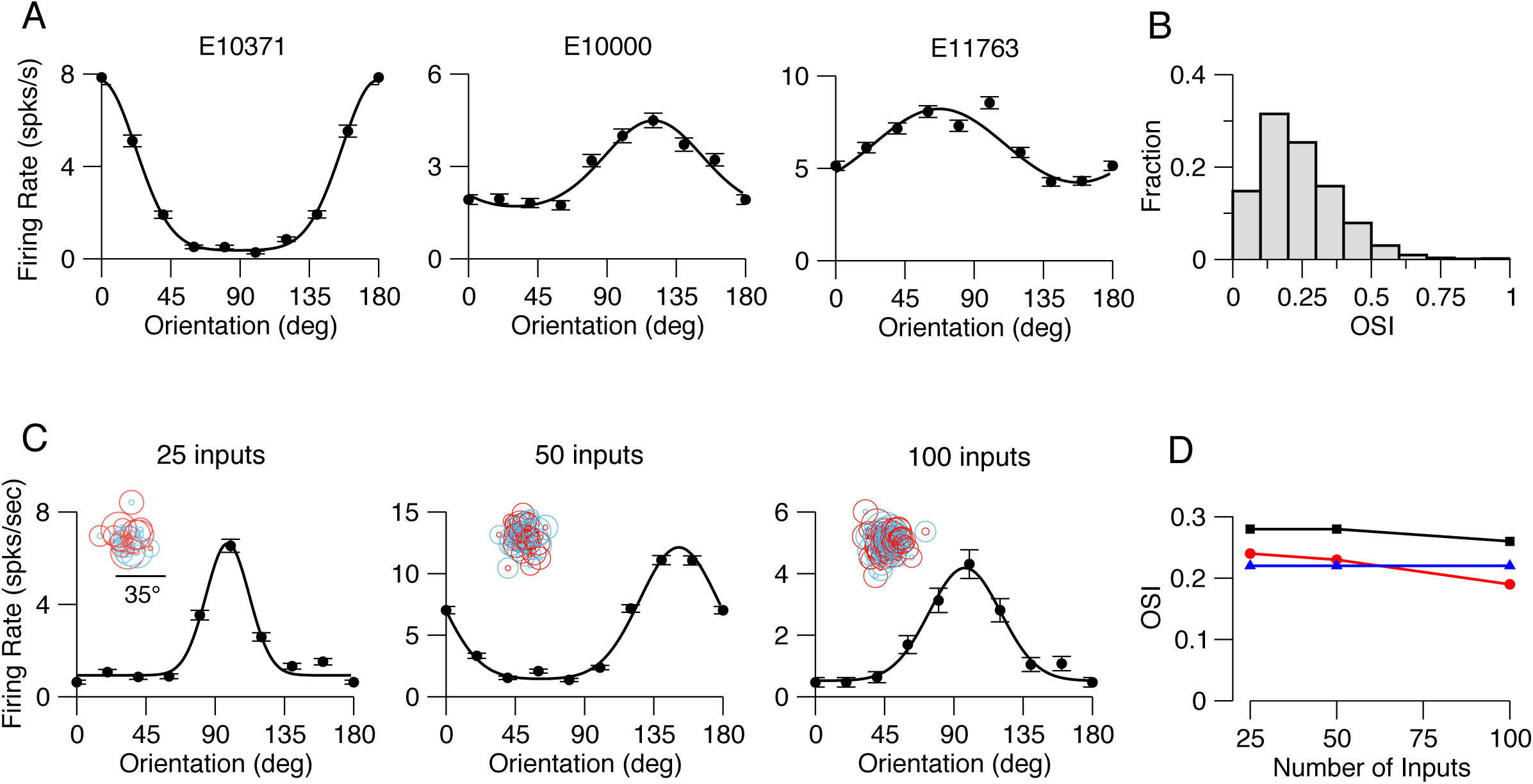
Orientation selectivity emerges in the mouse V1 model. **A.** Examples of tuning curves (peak firing rate) of three excitatory V1 neurons in the model. SF of the drifting grating is 0.03 cyc/deg. OSIs from left to right are: 0.62, 0.23, 0.15. **B.** Distribution of OSI (peak response) over all the neurons (neurons in the central part of the network; see Methods; n=5041). Mean OSI=0.24 (mean OSIs of the F0 and F1 components of the response are 0.29 and 0.19). **C.** Examples of tuning curves of excitatory neurons in networks with different average number of thalamic inputs per neuron. From left to right: OSI=0.47, 0.48, 0.49. **D.** Average OSIs vs. average number of thalamic inputs. Red: Peak spike response. Black: F1 component of the spike response.Blue: F1 component of the thalamic excitatory input.

Orientation selectivity emerges in our random connectivity model because of the spatial inhomogeneity in inputs to cortical neurons. In particular, the convergence of ON and OFF thalamic inputs onto model neurons are spatially offset from one another. The orientation of this offset may be related to the emergent orientation preference of neurons (Lien and Scanziani, 2013; Liu et al., 2010). To assess this relationship, we estimated the ON and OFF subfields of the thalamic inputs by presenting spots at different locations to the model network as in Lien and Scanziani (2013) (see Methods). The estimated ON and OFF subfields for 4 example neurons reveal different offsets. When ON and OFF subfields have large horizontal displacements (E14493, E14847) preference for the vertical orientation of the drifting grating at 0.03 cyc/deg tends to emerge whereas when ON and OFF subfields are vertically displaced preference for horizontal orientations tends to emerge (Fig. 3A, E14664). The offsets in ON and OFF subfields that emerge from the random connectivity model (Fig. 3B) are similar to those observed experimentally (Lien and Scanziani, 2013). When the ON/OFF offset is large there is a strong correspondence between the axis of the offset and the preferred orientation of the thalamic input (Fig. 3C). The ON and OFF displacement, however, is not the only factor that contributes to this orientation preference. The randomness in the feedforward connectivity generates ON and OFF subfields of the thalamic excitation that deviate from circularity. The shape of the subfields, and the interaction between the subfields, can create orientation preferences that deviate from that predicted from the offset of ON and OFF subfields (Fig. 3A, example 15022). In sum, the offset of ON and OFF subfields, their interaction, and their shape influence the emergent thalamic orientation selectivity. Because the thalamic input selectivity is directly related to the cortical output selectivity (Sup. Fig. 3C,D) these factors impact the emergent cortical orientation selectivity in the same way. The emergent orientation preference, however, is particularly sensitive to the spatial structure of the stimulus (Fig. 1).

**Figure 3:**
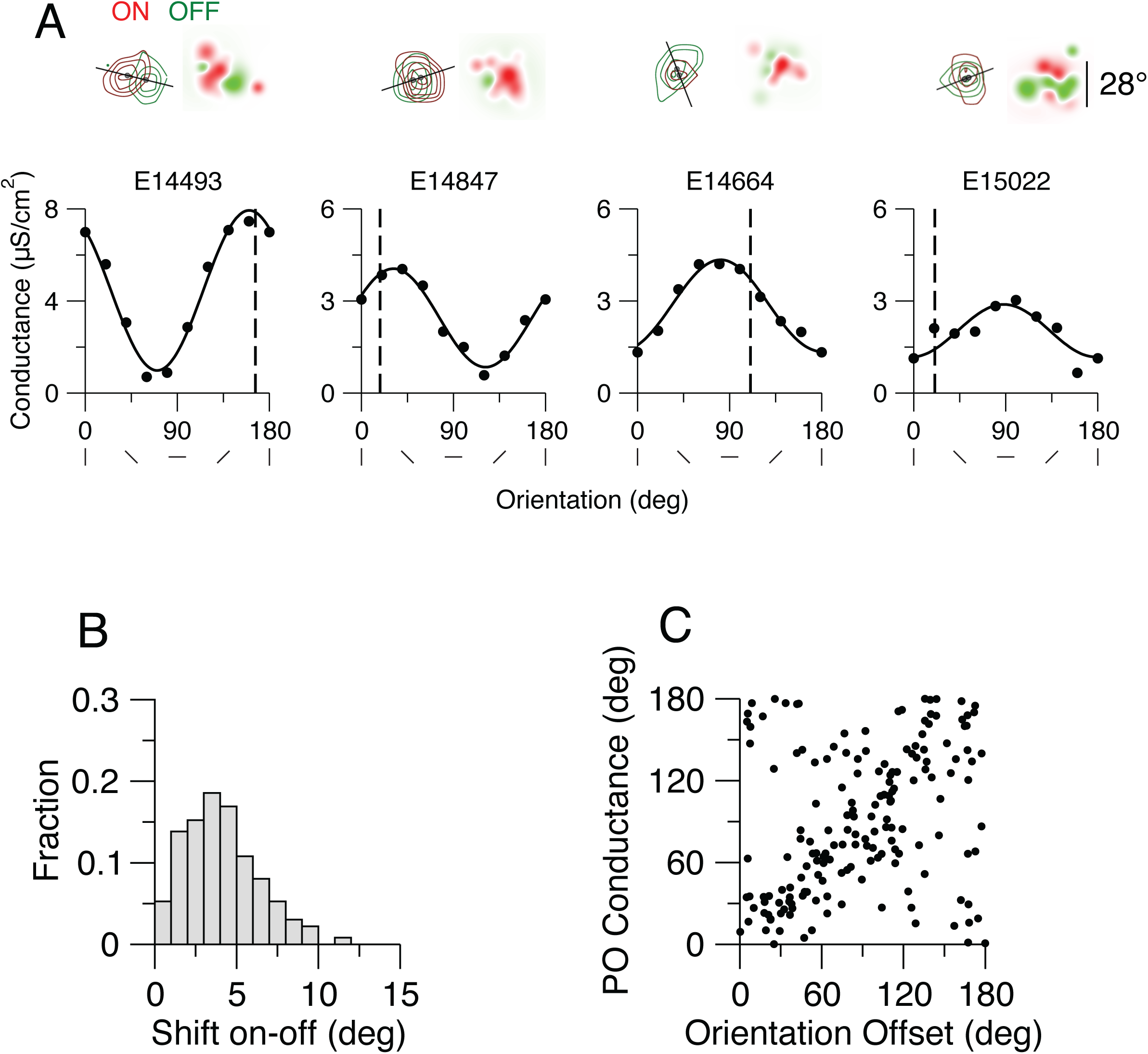
The contribution of the offset of ON and OFF subregions of the thalamic excitation to its orientation preference. The ON and OFF subfields of the thalamic inputs were estimated by presenting spots at different locations to the model network as in Lien and Scanziani (2013) (see Methods). **A.** Top panels: ON (red) and OFF (green) subfields of the thalamic excitation for four example neurons. Dark spots: Center of mass of the subfields. The solid line indicates the axis of the offset of the two centers of mass. Receptive fields based on the summed ON and OFF thalamic inputs are shown on the right. Bottom panels: Tuning curves of the thalamic excitation for these neurons. The SF of the drifting grating is 0.03 cyc/deg. Vertical dashed-line indicates the orientation of the offset axis (0° corresponds to an horizontal axis). Offset amplitude and orientation and preference of the thalamic excitation is: E14493: 11.4°; 166.1°, 160.3°. E14847: 4.7°, 18.2°, 31.1°. E14664: 3.9°, 111.4°, 80.7°. E15022: 2.8°, 20.6°, 88.0°. **B.** Offset distribution across neurons (n=361; neurons are at the center of the network, see Methods). Mean offset: 4.1**°. D.** Orientation preference of the thalamic input conductance (drifting grating with 0.03 cyc/deg) vs. orientation of the offset axis for all neurons with an offset larger than 4° (n=170). The circular correlation is: 0.24.

### Dependence of preferred orientation on spatial frequency in the model

We then characterized how much the properties of the neuronal responses vary with spatial frequency in the model. First we investigated how the population average peak response and OSI were affected when changing spatial frequency (SF, Sup. Fig. 3A). We found that although the mean population response was modulated by spatial frequency (maximal response for SF=0.035 cyc/deg) the overall selectivity of the population was less sensitive to SF (Sup. Fig. 3A,B). This mild effect across the population contrasts with the effect of spatial frequency changes on the preferred orientation of individual neurons. As we varied spatial frequency the preferred orientation of neurons often changed (top and bottom left panels in Fig. 4A; Fig. 4B,C, pink). We quantified this change by computing the circular correlation (CC, see Methods) of the preferred orientation at different spatial frequencies across neurons. This correlation was strong for nearby spatial frequencies whereas for spatial frequencies far apart it was (Fig. 4B,C). It declined from 0.71 for 0.04-0.03 cyc/deg to 0.00 for 0.04-0.01 cyc/deg (Fig 5, ΔCC=0.71). We found that this effect was robust to changes in the network size, the number of connections per neuron and the synaptic conductance strengths (Sup. Fig. 5B,D). We also found that it was qualitatively robust to changes in the spatial dispersion of the thalamic feedforward connections but that the decorrelation was weaker for smaller dispersions (Sup. Fig. 6).

**Figure 4:**
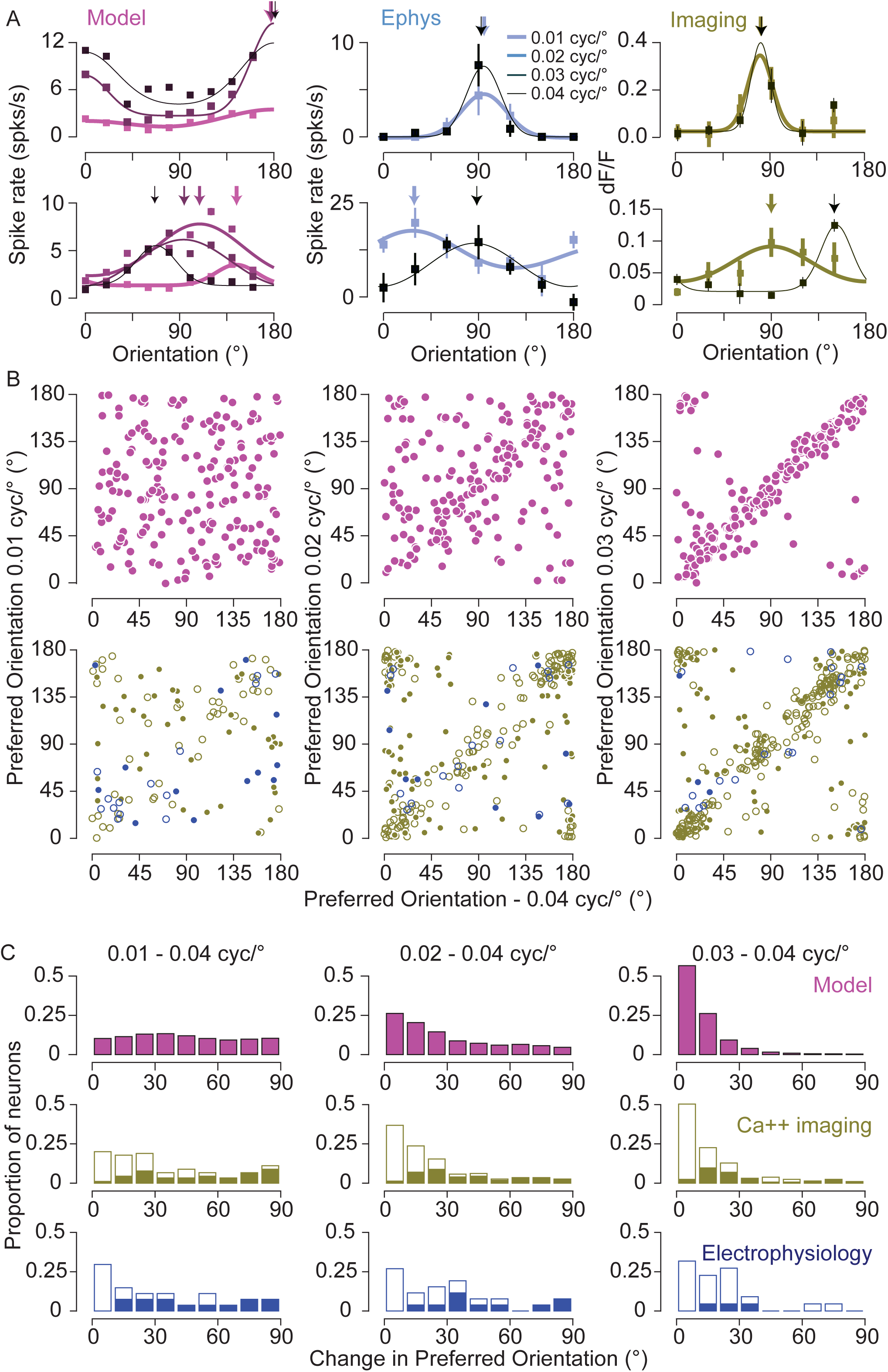
Spatial frequency and orientation selectivity in the model and mouse V1. **A.** Example orientation tuning curves based on spike rate are plotted for neurons in the spiking network model (left), electrophysiology (middle) and based on fluorescence changes from calcium imaging experiments (right). Orientation tuning curves are plotted for different spatial frequencies, from 0.01 to 0.04 cyc/deg, indicated by color. If the error bars are not visible, they are smaller than the symbol size. **B.** Top row: The relationship between preferred orientation in the model. Left: 0.04 cyc/deg and 0.01 cyc/deg. Middle: 0.04 cyc/deg and 0.02 cyc/deg. Right: 0.04 cyc/deg and 0.03 cyc/deg. Bottom row: The same for the calcium and electrophysiological records (green and blue symbols, respectively). The bootstrapped vector average is used as the estimate of the preferred orientation. For calcium and spiking data, statistically significant shifts in orientation preference are indicated by filled circles. Number of cells in the imaging data for comparison of 0.01 and 0.04 cyc/deg is 90, for comparison of 0.02 and 0.04 cyc/deg is 228 and for comparison of 0.03 and 0.04 cyc/deg is 288. Number of cells in the electrophysiological data for comparison of 0.01 and 0.04 cyc/deg is 32, for comparison of 0.02 and 0.04 cyc/deg is 28 and for comparison of 0.03 and 0.04 cyc/deg is 25. **C.** Histograms of the difference in orientation preference between 0.04 cyc/deg and 0.01 (left), 0.02 (middle) and 0.03 (right) cyc/deg. Filled bars for electrophysiology and calcium imaging data indicate statistically significant changes in orientation preference.

### Dependence of preferred orientation on spatial frequency in mouse V1

These theoretical results prompted us to determine whether spatial frequency has a similar effect on orientation preference in mouse V1. Varying spatial frequency yielded shifts in orientation preference for many, but not all, neurons when measured using intracellular, whole-cell, recordings (Fig. 4A, middle: top and bottom panels; Fig. 4C, blue panels). Changes in orientation preference were observed both at the level of spike rate and membrane potential (38 total cells) (Sup. Fig. 4). To gain access to this effect in large populations of V1 neurons we also examined it by measuring calcium responses using 2 photon microscopy (606 total cells) (Fig. 4A, left: top and bottom panels; Fig. 4B, C, green panels; Sup Fig. 7). As with our electrophysiological data we found a diversity of changes with spatial frequency: preference shifted dramatically for some neurons and not for others.

These differences in preferred orientation observed from our Ca++ responses could be due to noise in our measurements. To be included in our population analysis cells were required to have a minimum peak response of 8% at both frequencies. Using different thresholds to include cells yields similar declines in correlation when comparing orientation preference at 0.04 cyc/deg to 0.03 and 0.01 cyc/deg (8%: ΔCC=0.49, 10%: ΔCC=0.4, 12%: ΔCC=0.52). To address whether the observed effect was influenced by differences in response amplitude for different spatial frequencies we also restricted our analysis to neurons with differences in peak response amplitudes less than 10%. This also did not alter the decline in circular correlation (ΔCC = 0.49). In sum, orientation preference changed with spatial frequency in electrophysiology records as well as calcium imaging measurements.

We have found that both the model and actual mouse V1 neurons exhibit changes in orientation preference with spatial frequency in a similar fashion (Fig. 5). That is, for small frequency shifts the model and actual neurons have similar orientation preferences, as indicated by a high circular correlation, whereas large changes in spatial frequency cause substantial decreases in circular correlation. One notable discrepancy between the model and actual data is that nearby spatial frequencies have higher correlations for the model than for the data. A factor that contributes to this discrepancy is the amount of data collected in the model records relative to the physiological records. When we limit the records from which the model data are based to 20 seconds, instead of 80 seconds, ΔCC declines from 0.71 to 0.58. An additional factor we considered is the nature of the thalamocortical input. Orientation selectivity does exist in mouse thalamic neurons (Piscopo et al., 2013; Scholl et al., 2013a; Zhao et al., 2013), so we also explored the impact of elongated thalamic receptive fields on the properties of the cortical model (Sup. Fig. 8). This impact was modest, slightly altering the dependence of orientation preference on spatial frequency (Fig. 5, elongated thalamic receptive field model, ΔCC=0.73, Supp. Fig. 8 F,G), while increasing the overall orientation selectivity(mean OSI=0.32 vs 0.23 for circular thalamic receptive field), Sup. Fig. 8B).

**Figure 5:**
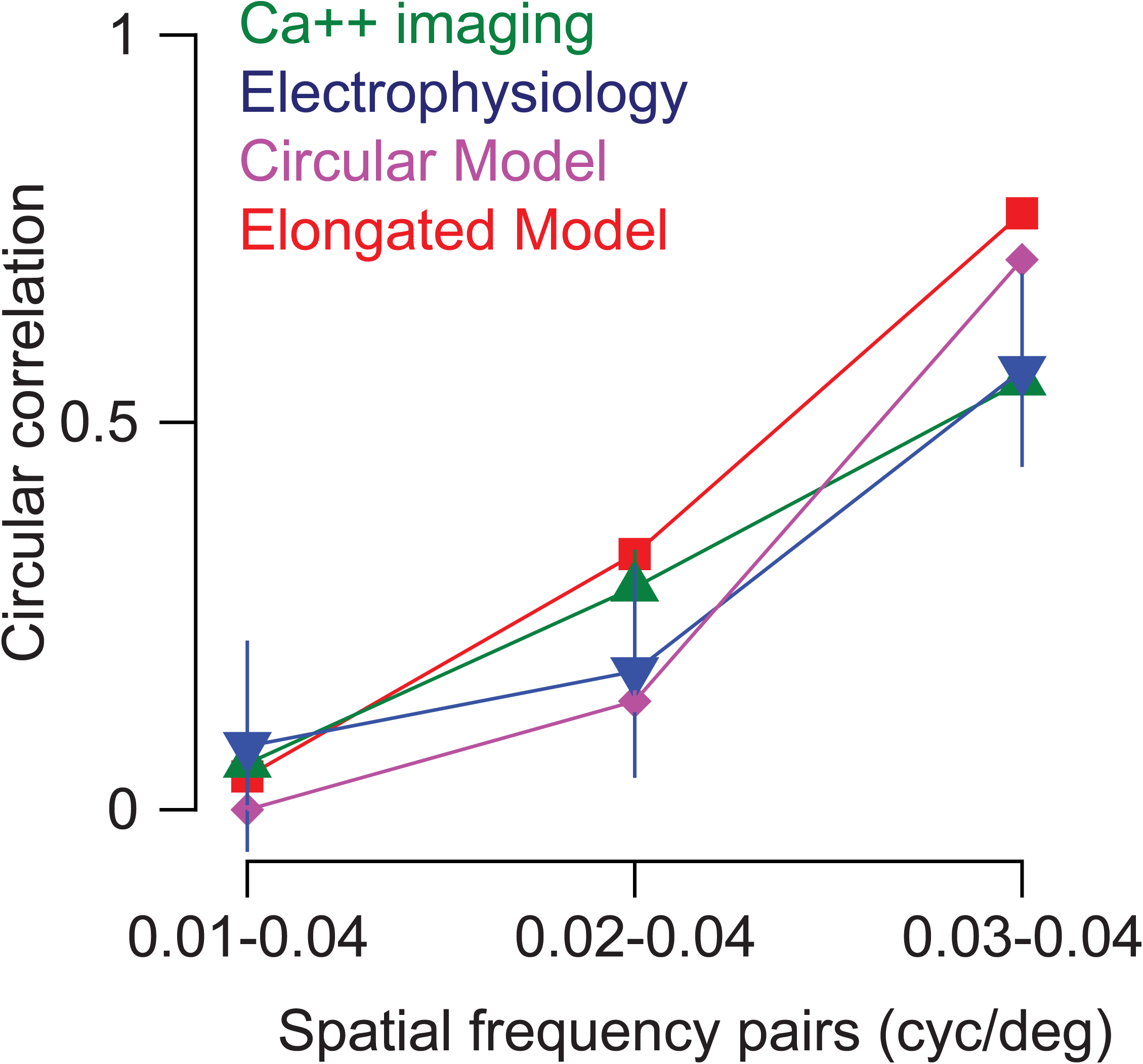
Comparison between model and experimental results. Graph indicates the observed circular correlation between preferred orientations of single neurons at two spatial frequencies. The pairs of spatial frequencies being compared are indicated on the X axis. Green: Calcium imaging. Blue: Electrophysiology,.Purple: Model with circular thalamic receptive fields (same as in Fig. 4), Red: Model with elongated thalamic receptive fields (see text and Sup. Fig. 9). Error bars are bootstrapped confidence intervals on the circular correlation.

### 2-D spatial frequency filters of neurons in mouse V1 are non-separable

The observed dependence of orientation preference on spatial frequency indicates that in mouse V1, neuron receptive fields are not simple orientation detectors. Instead they may be measuring components of the visual scene that are better characterized by a conjunction of 2-dimensional spatial frequency filters. We therefore measured responses of V1 neurons while varying vertical and horizontal spatial frequency components (Ringach et al., 2016) (24 cells, Hartley gratings, see Methods; Fig. 6A). Neurons whose orientation selectivity is invariant to spatial frequency, would exhibit preference profiles for which angle (orientation) does not change with the distance from the origin (spatial frequency). As before different neurons revealed a diversity of behaviors (similar to kernels shown in Ringach et al., 2016), from invariance (Fig. 6A, left) to systematic change in selectivity with spatial frequency (Fig. 6A, middle). We also recorded from a small number of inhibitory neurons with broad selectivity for orientation and spatial frequency (Niell and Stryker, 2008) Fig. 6A, right). Measures of orientation preference based on the Hartley stimulus qualitatively agree with those made by measuring orientation tuning curves at different spatial frequencies (compare top and bottom panels in Fig. 6A). This indicates that many V1 neurons are better characterized as containing receptive fields that are a conjunction of horizontal and vertical spatial frequency filters instead of invariant selectivity for orientation. We performed a comparable analysis in our V1 network model (see Methods) and found a similar behavior (Fig. 6B; Sup. Fig. 9).

**Figure 6:**
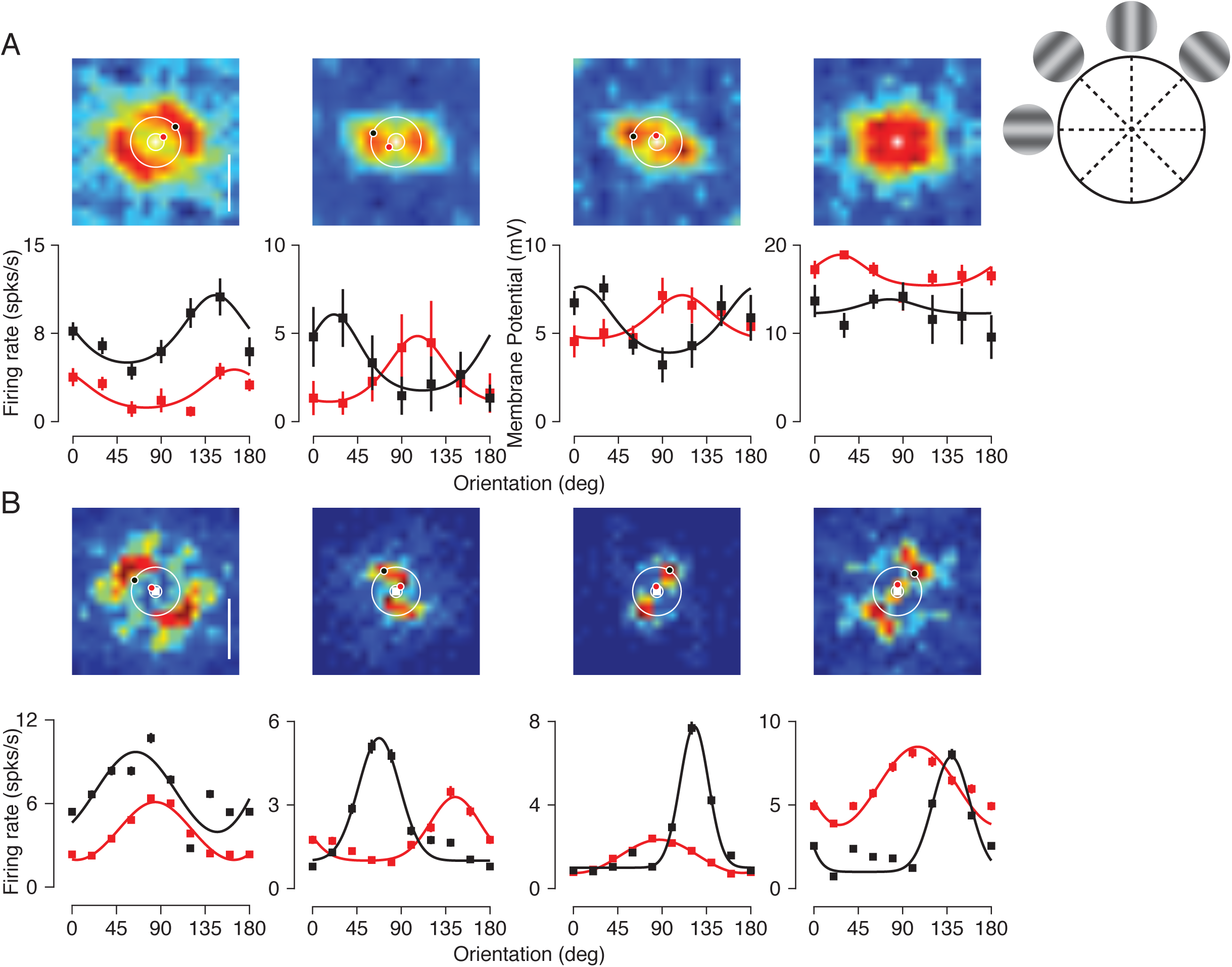
Neuron receptive fields in the frequency domain are intricate. **A.** Mean membrane potential responses to Hartley stimuli (see Methods) are plotted for combinations of horizontal and vertical spatial frequencies (top row). Circles indicate stimulus combinations corresponding to oriented gratings at fixed spatial frequencies. Each panel corresponds to a different example cell. Orientation tuning curves for drifting gratings at 0.014 cyc/deg and 0.044 cyc/deg are shown for these four neurons (bottom row). **B.** Example frequency receptive fields for four neurons in the model. Orientation tuning curves at 0.01 cyc/deg and 0.04 cyc/deg are shown for these neurons (bottom row) based on responses to drifting gratings.

## Discussion

We have presented a network model for rodent V1 that demonstrates that orientation selectivity can emerge from random connectivity even if LGN cells are not selective. It makes the specific prediction that this selectivity should be sensitive to spatial form. Testing that prediction in mouse visual cortex we found a similar effect. Using a model that receives thalamic inputs which exhibited some orientation selectivity increased the degree of cortical orientation selectivity yielding distributions of OSI closer to experimental estimates. This model also exhibited a similar dependence of orientation preference on spatial frequency.

In our models there is a strong overlap of the ON and OFF subregions of the thalamic inputs as seen in experiments (Li et al., 2013; Lien and Scanziani, 2013; Liu et al., 2010). When the offset between the centers of the ON and OFF subfields is large, the orientation of this offset can be predictive of the orientation preference of the neuronal response. Nevertheless, even when this offset is large, the orientation preference can change substantially with spatial frequency. In our model the orientation of the offset and the orientation preference of the neuronal response are strongly correlated for intermediate spatial frequency only (Sup. Fig. 10).

Quantitatively, the decorrelation of preferred orientation with spatial frequency is somewhat weaker in experiments when compared to our models. One source of this discrepancy is related to the amount of data collected for the model and the experiments. When records for the model are limited to 20 seconds, the model ΔCC was 0.59, close to the experimental value of 0.49. Another possible source for this difference is that we did not incorporate any feature specific component in the connectivity even though this has been shown to be present in mouse V1 after the critical period (Ko et al., 2013; Ko et al., 2011; Lee et al., 2016b).

We have demonstrated that V1 neurons’ receptive fields are surprisingly intricate (Fig. 6, Supp. Fig. 9). This complexity stands in contrast to the V1 receptive fields in cats (Hammond and Pomfrett, 1990; Jones et al., 1987; Webster and De Valois, 1985) and primates (De Valois et al., 1982), where orientation preference is represented in a separable manner from spatial form. A similar dependence in the mouse V1 was reported in a study based on calcium imaging (Ayzenshtat et al., 2016). There it was demonstrated that a reduction in spatial frequency by one octave causes a mean shift in preferred orientation by 22.1°, comparable to our own estimates of the change in orientation when shifting from 0.04 to 0.02 cyc/deg (model: mean ΔPO = 29.8°, ephys: ΔPO =30.2°, ca++: ΔPO =22.2°). They proposed that the dependence could arise from separable selectivity in frequency domain. We demonstrate here that while some V1 neurons do have separable frequency domain receptive fields, V1 receptive fields exhibit diverse dependencies that yield spatial frequency invariant orientation preferences (Fig. 6, first column) or spatial frequency dependent orientation preferences (Fig. 6, second column).

Such receptive field complexity likely has an impact on connectivity patterns within V1. In primates and carnivores where preferred orientations are similar for different spatial frequencies, neurons with similar orientation preferences are much more likely to be connected (Bosking et al., 1997; Wilson et al., 2016). In mice, neurons with similar orientation preference have been reported to be somewhat more likely to be connected (Ko et al., 2013; Ko et al., 2011). However, in these experiments difference in preferred orientation was measured at only one spatial frequency (0.045 cyc/deg). As we have shown this difference varies with spatial frequency and the connectivity is likely to depend on the similarity in response at all spatial frequencies. Indeed, correlation in the response to natural stimuli was found to be a stronger factor than orientation preference at one spatial frequency in determining connection probability (Cossell et al., 2015; Ko et al., 2013).

The intricate receptive field profiles described here are akin to those observed in primary auditory cortex. Auditory cortex neurons are sensitive to the combination of many auditory cues (Wang et al., 2005), which may comprise a synthesis sufficient to detect auditory objects (Bar-Yosef and Nelken, 2007). The frequency domain receptive field profiles observed in mouse V1 neurons may therefore reflect a similar progression toward a representation for objects using a random connectivity scheme that occurs as information flows through the visual pathway.

To conclude, our investigation demonstrates that random connectivity can be the dominant component accounting for emergent properties such as orientation selectivity. An important advantage of random wiring schemes is that they occur naturally, following the broader patterns of retinotopy that are formed by biochemical gradients. This natural emergence may thus reflect a wiring strategy that allows for selectivity without the cost associated with constructing specific afferent wiring connections.

## Methods

Procedures for two-photon imaging and physiology were based on those previously described (Scholl et al., 2015; Scholl et al., 2013b). Animals were anesthetized for the duration of the experiments and stimuli were presented on calibrated CRT monitors. Analyses of physiological data were performed using routines in Matlab that have previously been employed (Scholl et al., 2013b). The computational model is composed of two networks. One represents LGN while the other represents layer 4 and layer 2/3. Neurons of the cortical network are described in terms of conductance-based models (Hansel and van Vreeswijk, 2012; Wang and Buzsaki, 1996). Details describing both the experimental and computational approaches are in the Online Methods Section.

## Acknowledgements

We are grateful to Ran Darshan, Carole Levenes and Gianluigi Mongillo for helpful discussions and comments and Baowang Li for conducting some of the electrophysiology experiments. Supported by a CRCNS grant (ANR: 14-NEUC-0001-01, NIH: EY-020592), ANR-13-BSV4-0014-02 (CvV and DH), CONICET grant PIP 112 201301 00256 (GM). Part of the modeling work was performed in the framework of the France-Israel Laboratory of Neuroscience (DH and CvV). JJP is a Howard Hughes Medical Institute International Student Research Fellow.

## Online Methods and Supplementary Materials

### Detailed experimental methods

#### Physiology

Procedures for two-photon imaging and physiology were based on those previously described (Scholl et al., 2015; Scholl et al., 2016). Experiments were conducted using normal, adult male and female animals (n =33, P34 - P60). Mice were anesthetized with intraperitoneal injections of 1000 mg/kg urethane and 10 mg/kg chlorprothixene. Brain edema was prevented by intraperitoneal injection of up to 10 mg/kg dexamethasone. Animals were warmed with a thermostatically controlled heat lamp to maintain body temperature at 37° C. A tracheotomy was performed and the head was placed in a mouse adaptor (Stoelting). A craniotomy and duratomy were performed over visual cortex. Eyes were kept moist with a thin layer of silicone oil. Primary visual cortex was located and mapped by multi-unit extracellular recordings with tungsten electrodes (1 mΩ, Micro Probes). The V1/V2 boundary was identified by the characteristic gradient in receptive locations (Drager, 1975; Metin et al., 1988). Eye drift under urethane anesthesia is typically small and results in a change in eye position of less than 2 degrees per hour(Sarnaik et al., 2014).

#### Dye Loading and In vivo Two-Photon Microscopy

Bulk loading of a calcium sensitive dye under continuous visual guidance followed previous protocols in V1 (Golshani and Portera-Cailliau, 2008; Kerr et al., 2005; Mrsic-Flogel et al., 2007; Ohki et al., 2005; Stosiek et al., 2003). Dye solution contained 0.8 mM Oregon Green 488 BAPTA-1 AM (OGB-1 AM, Invitrogen) dissolved in DMSO (Sigma-Aldrich) with 20% pluronic acid (Sigma-Aldrich) and mixed in a salt solution (150 mM NaCl, 2.5 mM KCl, 10 mM HEPES, pH 7.4, all Sigma-Aldrich). 40-80 µM Alexa Fluor 594 (Invitrogen) was also included for visualization during and immediately after loading. Patch pipettes (tip diameter 2-5 µm, King Precision Glass) containing this solution were inserted into the cortex to a depth of 250-400 µm below the surface with 1.5% agarose (in saline) placed on top the brain. The solution was carefully pressure injected (100-350 mbar) over 10-15 minutes to cause the least amount of tissue damage. OGB-1-AM is only weakly fluorescent before being internalized, so the amount of dye injected was inferred through the red dye. To ensure full loading we waited 1 hour before adding a glass coverslip for imaging. Metal springs were fastened on the attached head plate to place pressure on the glass coverslip and reduce brain pulsations. Fluctuations in calcium fluorescence were collected with a custom-built two-photon resonant mirror scanning microscope (Scholl et al., 2015) and a mode-locked (925 nm) Chameleon Ultra Ti:Sapphire laser (Coherent). Excitation light was focused by a 16X or 40x water objective (0.8 numerical aperture, Nikon). Images were obtained with custom software (Labview, National Instruments). A square region of cortex 300 µm wide was imaged at 256×455 pixels. In all experiments, multiple focal planes, separated by 20-25 µm, were used to collect data, starting around 150 µm below the cortical surface. Before each experiment neuron drift was measured over a 2-3 min period. If drift occurred then the glass coverslip and agarose were readjusted to stabilize the brain during stimulus protocol (7-20 minutes each focal plane).

#### Stimuli

Visual stimuli were generated by a Macintosh computer (Apple) using the Psychophysics Toolbox (Brainard, 1997; Pelli, 1997) for Matlab (Mathworks). Gratings were presented using a Sony video monitor (GDM-F520) placed 25 cm from the animal’s eyes. The video monitors had a non-interlaced refresh rate of 100Hz,a spatial resolution of 1024×768 pixels, which subtended 40×30 cm, and a mean luminance of 40 cd/cm^2^. Drifting gratings (38 deg diameter for imaging, variable diameter for electrophysiology, 0.01-0.04 spatial frequency, 100% contrast, 2 Hz temporal frequency) were presented for 2-3 sec. Each stimulus was followed by a 3 sec blank (mean luminance) period in the imaging protocol. Spontaneous activity was measured during blank (mean luminance) periods interleaved with drifting grating stimuli, all presented in a pseudorandom sequence. Direction presented ranged from 0-330 deg. Different spatial frequencies used were either presented individually in separate blocks or interleaved within the same block. Hartley stimuli were presented for each spatial frequency combination for 250 ms (Malone and Ringach, 2008; Ringach et al., 2016). For each spatial frequency combination four phases were presented and the response to these phases were averaged. These were repeated 5-30 times per cell. During imaging sessions, each stimulation protocol was repeated 7-10 times at each focal plane. The microscope objective and photomultiplier tubes were shielded from stray light and the video monitors.

#### Two-photon Calcium Imaging Analysis

Images were analyzed with custom Matlab software (Mathworks). Cells were identified by hand from structure images based on size, shape, and brightness. Cell masks were generated automatically following previous methods(Nauhaus et al., 2012). Glia were easily avoided due to their different morphology from both OGB-1 AM filled neurons. Time courses for individual neurons were extracted by summing pixel intensity values within cell masks in each frame. Responses (*F*_*t*_) to each stimulus presentation were normalized by the response to the gray screen (*F*_0_) immediately before the stimulus came on:

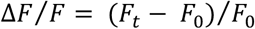

For each stimulus, the mean change in fluorescence Δ*F/F* was calculated in a 0.66 sec window of the response, identified by averaging responses to all stimuli and detecting the global peak. Visually responsive cells were identified if at least one orientation evoked a response with:

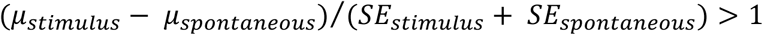

where *µ*_*stimulus*_ refers to the mean stimulus evoked response, *µ*_*spontaneous*_ refers to the mean spontaneous activity, *SE*_*stimulus*_ is the stimulus evoked response standard error, and *SE*_*spontaneous*_ spontaneous activity standard error. Additionally, identified responses were required to have distinct different trial-to-trial fluorescence time courses, so as to not be scaled versions of neuropil activity. The maximum peak response for each cell was also required to have a response amplitude greater than 0.08. Mean changes in fluorescence from visually responsive neurons were used to generate tuning curves for orientation selectivity.

#### Electrophysiology Analysis

Spiking responses for each stimulus were cycled-averaged across trials after removing the first cycle. The Fourier transform of mean cycle-average responses was used to calculate the mean (F0) and modulation amplitude (F1) of each cycle-averaged response, after mean spontaneous activity was subtracted. The subthreshold membrane potential responses were also similarly computed after median filtering the voltage traces to remove spikes. Peak responses were defined as the sum of the mean and modulation (F0 + F1).

Peak responses per trial across each condition for neuronal responses measured using electrophysiology and imaging were bootstrapped to compute the vector average orientation. This was used as the preferred orientation for the neuron. For electrophysiology, cells were only included in the analysis, if the bootstrapped confidence intervals on mean of the maximum amplitude spiking response did not include zero. A double Gaussian curve was fit to the responses for characterizing orientation tuning (Carandini and Ferster, 2000):

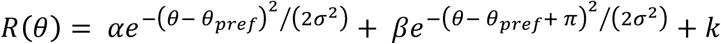

Here *R*(*θ*) is the response of the neuron to different orientations (*θ*), σ is the width of the tuning curve, *k* is the mean activity, α and β are peak amplitudes, and *θ*_*pref*_ is the orientation preference. Gaussian fits were used only for qualitative description of the tuning. The actual fit parameters have not been used in the analysis. The orientation selectivity index was also computed (Ringach et al., 2002; Tan et al., 2011):

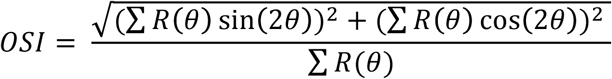

The circular correlation (cc) between the preferred orientations (PO) is defined as:

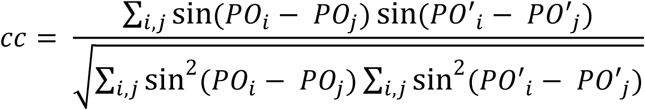

where PO_i_ is the preferred orientation of neuron i for one spatial frequency and PO’_i_ is the preferred orientation of the same neuron for another spatial frequency. This number is always in the range [-1:1], reaching 1 for perfect linear correlation between the preferred orientations in the two conditions.

To determine if the difference in the preferred orientations computed at different spatial frequencies was statistically significant, we generated bootstrapped confidence intervals on the both the preferred orientations being compared. The difference was considered significant if these intervals did not overlap.

### The computational model of mouse V1

The model is composed of two networks. One represents LGN and has N_L_ neurons. The second network represents layer 4 and layer 2/3 in mouse V1. For simplicity these two layers are collapsed into one single network, with N_E_ excitatory and N_I_ inhibitory neurons. In both networks the neurons are arranged on a square grid and the position (x_iA_,y_iA_), where (i,A) denotes the neuron i=1,…,N_A_ of population A=E,I,L. The position of neuron (i,A) is given by 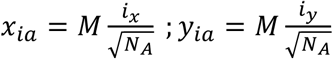 where M is the size of the network (2mm), 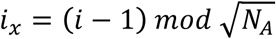 and 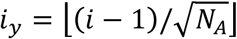. Here is the largest integer equal to or smaller than x. All N_A_ are square integers so that i_x_ and i_y_ are integers between 0 and 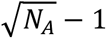. Unless said otherwise we take N_E_=32400, N_I_=8100, N_L_=25600.

#### Cortical neurons

They are described in terms of conductance-based models. The membrane potential of neuron (i,A), A=E,I, evolves in time according to

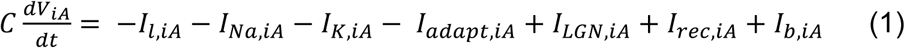

where C is the membrane capacitance, *I*_*l,iA*_, is the leak current, and I_Na,iA_, I_K,iA_ are the intrinsic sodium and potassium currents that shape the action potentials and I_adapt,iA_ is an adaptation potassium current which included in E neurons, only. The dynamics of these currents are as in (Hansel and van Vreeswijk, 2012). The current I_LGN,iA_ describes the input from LGN, I_rec,iA_ is the recurrent input from other cortical neurons and I_back, iA_ represents a background input from other cortical regions not explicitly included in the model.

#### LGN neurons

LGN cells are modeled as Poisson neurons with time varying rates that depend on the visual stimulus. Neuron (i,L) responds to a luminosity field L(x,y,t) with an instantaneous firing rate

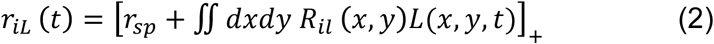

where r_sp_ is the spontaneous firing rate of the neuron, assumed to be the same for all LGN cells, R_iL_(x,y) is its receptive field and [x]_+_=x for x >0, [x]_+_=0 for x<0.. The luminosity field of a sinusoidal drifting grating with orientation, θ, spatial wavelength, λ, and temporal frequency, ω, is

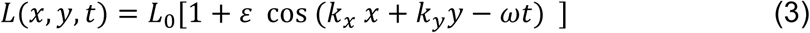

where *L*_0_is the average luminosity, ε is the contrast, and the wave-vector of the grating is:***k*** = (*k*_*x*_,*k*_*y*_) = (*k* cos θ,*k* sin θ) with *k* = 2 π/ λ. The parameters used in our simulations are listed in Tables 1 and 2.

**Table 1:** Default parameters of the computational model of mouse V1 (Layer 4 and stimulus).

|  |  |
| --- | --- |
| $N_E$ | 32400 |
| $N_I$ | 8100 |
| $K_{AB} (A,B=E,I)$ | 500 |
| $K_{EL}$ | 25 |
| $K_{IL}$ | 100 |
| $g_{EE}$ | 0.011 ms mS/cm<sup>2</sup> |
| $g_{IE}$ | 0.067 ms mS/cm<sup>2</sup> |
| $g_{EI}$ | 0.029 ms mS/cm<sup>2</sup> |
| $g_{II}$ | 0.098 ms mS/cm<sup>2</sup> |
| $g_{EL}$ | 0.04 ms mS/cm<sup>2</sup> |
| $g_{iL}$ | 0.02 ms mS/cm<sup>2</sup> |
| $\tau_{AB} (A=E,I ; B=E,I,L)$ | 3 ms |
| $\sigma_{AB} (A,B=E,I)$ | 200 μm |
| $\sigma_{AL} (A=E,I)$ | 100 μm |
| $V_E$ | 0 mV |
| $V_I$ | -80 mV |
| $V_L$ | -65 mV |
| $\rho$ | 0.5 |
| $K_b$ | 500 |
| $g_b$ | 0 |
| $r_0$ | 1 Hz |
| $L_0$ | 0.075 |
| $\varepsilon$ | 1 |

**Table 2:** Parameters of the computational model of mouse V1: default values for the LGN cells.

|  |  |
| --- | --- |
| $N_L$ | 25600 |
| $r_{sp}$ | 1 Hz |
| $\gamma$ | 1 |
| $\sigma_{cxm} (*)$ | 1.3 |
| $\sigma_{sxm} (*)$ | 2.7 |
| $\beta_m$ | 0.05 |
| $\sigma_{cxs} (*)$ | 0.2 |
| $\sigma_{sxs} (*)$ | 0.2 |
| $\beta_s$ | 0.45 |

The receptive field of neuron (i,L) has the form

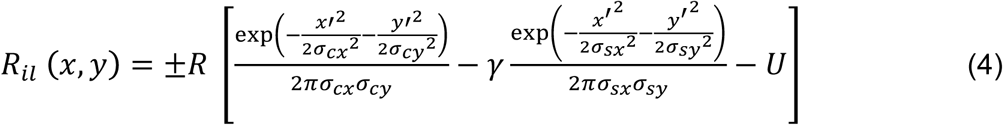

where 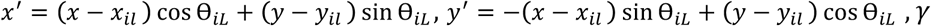 is a parameter that controls the relative weights of the two subfields, U is a constant such that 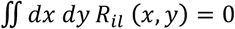 and R is a constant (1 Hz). The long and short axis of the center (resp. surround) region are denoted here by *σ*_*cx*_ and *σ*_*cy*_ (resp. *σ*_*sx*_ and *σ*_*sy*_). The global sign is +1 if the receptive field is ON center and −1 if it is OFF center. We take this sign at random with equal probability to be +1 or −1.

In all simulations except those in Supp. Fig. 8 we assume circular receptive fields for both center and surround subfields. In the simulations described in Supp Fig. 8 surrounds are circular but centers are elongated. We use the following parametrization: 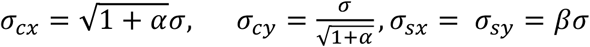 with 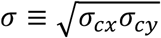. Therefore, *α* =0 corresponds to a circular center and surround subfields. In this case the LGN cell is not selective to orientation. The degree of selectivity increases with α.

The response of the LGN cells to a drifting grating can then be calculated based on

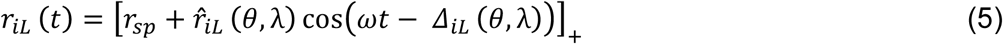

where

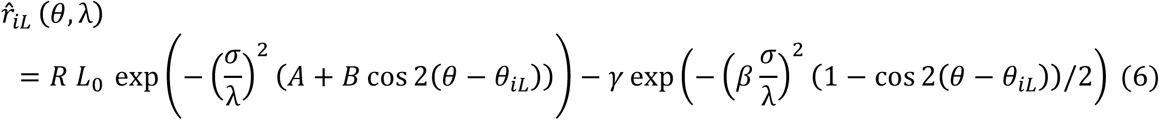

with *A* = (*α*^2^ +1)/2 and *B* = (*α*^2^ −1)/2.

The phase is: 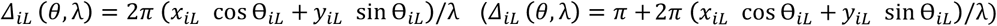 for an ON (OFF) cell.

#### Thalamo-cortical and recurrent connectivity

The connectivity between model LGN and cortex is random and does not depend on the functional properties of the cells. The probability that cortical neuron (i,A) is connected to LGN cell (j,L) is

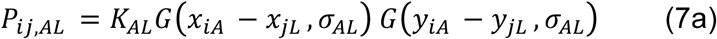

Where *K*_*AL*_ is the mean number of LGN inputs received by a cortical cell in population A. and

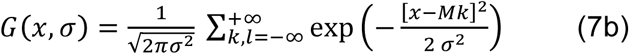

is the periodic Gaussian with variance *σ*^2^. The recurrent interactions in the cortical network are also random and non specific. The probability of connection between neuron (j,B) and (i,A) (A=E,I; B=E,I) is

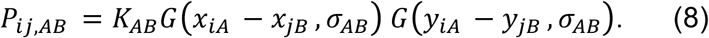

#### The feedforward and recurrent synaptic currents

Thalamo-cortical synapses on cortical population A are all excitatory, have a reversal potential *V*_*E*_, a strength *g*_*AL*_ and a synaptic time constant *τ*_*L*_. The thalamo-cortical current, *I*_*LGN,iA*_, in neuron (i,A) is

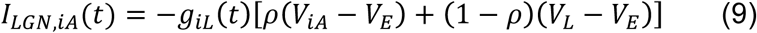

with: 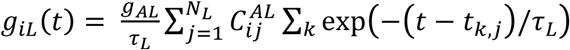, where *C*^*AL*^ is the *N*_*A*_ *X N*_*L*_ connectivity matrix of the thalamo-cortical projections (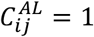 if there is a connection from neuron (j,L) to neuron (i,A); 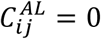 otherwise), and *t*_*k,j*_ is the time of the k-th spike generated by neuron (j,L). The sum over *k* is over all the spikes with *t*_*k,j*_ < *t*.

The total recurrent current into neuron (I,A) is *I*_*rec,iA*_ = *I*_*iA,E*_+*I*_*iA,I*_ where

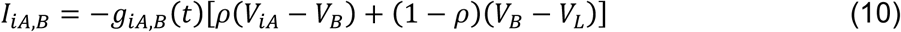

with 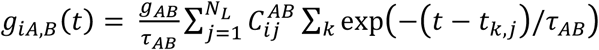.

Finally, the background current in Eq. (1) is modeled as

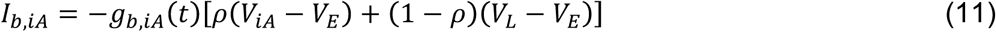

where *g*_*b,iA*_ (t) is a random Gaussian variable with mean *I*_*b*_*g* _*b*_*r*_0_ and variance 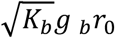. This represents the effect of *K*_*b*_ uncorrelated Poisson inputs, each of synaptic strength *g* _*b*_.

Note that in Eqs. (9,10) the right hand-sides comprise two contributions. The first is proportional to the driving force *V*_*iA*_ - *V*_*B*_. Thus it modifies the input conductance of the neuron. This contrasts with the second contribution which does not depend on the membrane potential of the post-synaptic cell. We adopted this description to incorporate in a simplified manner the fact that the change in input conductance induced by a synapse depends on its location on the dendritic tree. Proximal synapses which substantially affect the neuron’s input conductance are represented by the first contribution. The second contribution accounts for the synapses which are distal and which affect the input conductance of the neuron less (see also Hansel and van Vreeswijk, 2012).

#### Numerical procedures and analysis

Numerical simulations were performed using a 4th-order Runge-Kutta scheme to integrate the neuronal dynamics (Press, 1992). The synaptic interactions and the noise were treated at first order. The time step is *δt* = 0.05*ms*.

For each cortical neuron the mean firing rate, *F*_0_(*θ*_*k*_), and firing rate temporal modulation (first Fourier component of the response) *F*_1_(*θ*_*k*_) were estimated for each orientation, *θ*_*k*_(*k -* 1)20°, *k* = 1,..,9, by averaging the response upon 40s of stimulation, unless specified otherwise. We then computed the orientation averaged responses

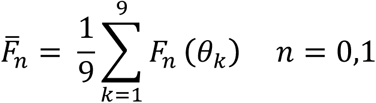

and the complex numbers

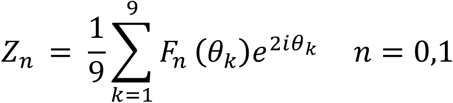

The Orientation Selectivity Index (OSI) and the Preferred Orientation (PO) of the peak response is then estimated from

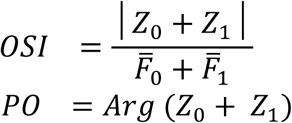

The OSI is 0 if the response has no tuning and 1 if the neuron responds at only one orientation. These definitions for the OSI and PO are equivalent to those used in the analysis of the experimental data (see above).

The definition of correlation coefficient is same as described above.

We also fit the tuning curves of the mean, *F*_0_(*θ*), and temporal modulation, *F*_1_(*θ*), of the spike to periodic Gaussian functions

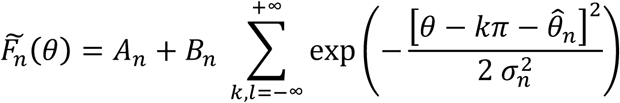

with n=0,1. We estimated the parameters 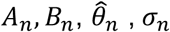, for each neuron by minimizing the quadratic error: 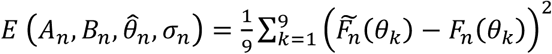

#### Robustness of the results

To check that a time step, *δt* = 0.05*ms*, was sufficiently small, we also performed several simulations with *δt* = 0.025*ms*. To verify that our results were also robust to changes in system size we performed several simulations on networks with N_E_=78560, N_I_=19600, N_L_=40000, keeping the average number of connections into E and I cells the same.

#### Structure of the ON and OFF subfield of the thalamic input

We characterized the thalamo-cortical input in the model by performing simulations with a protocol similar to the one in the experiments of Lien & Scanziani (2013). The stimuli used to map the receptive fields were Gaussian spots with a standard deviation of 5.6 degrees. The spots were presented in one of 64 locations arranged regularly in a square of 8×8 in the center of the network. The distance between the centers of adjacent spots was 7°. In order to characterize both ON and OFF receptive fields the stimuli were either brighter or dimmer than the background illumination. Each stimulus was presented during 1sec. During that time we evaluated the average of the conductance of the thalamic to each cortical neuron. We checked that the results were robust with respect to longer simulation times. The intensity of the stimulus (with respect to the background value) at the center of the Gaussian was l_0_=±0.075. After performing the simulations, the centers of the ON and OFF subfields were estimated by evaluating their center of mass: <**r**> = Σ_i_ f_i_ **r**_i_ /Σ_i_ f_i_, where f_i_ is the average thalamic input for a stimulus at position is **r**_i_. In order to reduce the noise level we performed the sum only over the locations for which the average input is larger or equal than 30% of the maximal average input.

Let us note that this way of estimating the center of the fields is only valid for cortical neurons whose feedforward inputs do not come from the border of the LGN network. Otherwise, because of the periodic boundary conditions of the LGN receptive fields, the linear estimation could combine inputs from opposite sides of the visual field. As the feedforward connectivity profile is topographically organized, neurons in the center of the cortex receive inputs from neurons in the center of the LGN. Therefore, boundary effects can be avoided by evaluating the center of mass only for neurons in the central part of the cortical network. In particular all the statistics of the ON and OFF subfields were estimated from neurons the square region of 14°×14° at the center of the network (361 neurons).

#### Parameters of the computational model

The cortical network is assumed to have a size of 2mm × 2mm representing 140°× 140° in the visual field(Kalatsky and Stryker, 2003).

The synaptic dispersion of the recurrent connectivity is taken to be 200 µm, consistently with values reported in Reyes & Sakmann, 1999 (Reyes and Sakmann, 1999). Unless indicated otherwise, the dispersion of the feed-forward connectivity was 100 µm.

The synaptic efficacies were as in Table 1. With these parameter values post-synaptic potentials have peak size is 0.5 mV (E->E interaction), −0.3 mV (I->E), 2.7 mV (E->I), - 0.9 mV (I->I), 0.9 mV (LGN->E), 0.8 mV (LGN->I). See Supplementary Figure 1a.

We introduced heterogeneity in the parameters *σ*_*cx*_, *σ*_*sx*_, *α, β*. For each thalamic neuron these parameters were chosen from a log-normal distribution

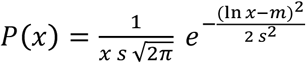

where the parameters *m* and *s* are given by *σ*_*cxm*_, *σ*_*cxs*_, *σ*_*sxm*_, *σ*_*sxs*_, *α*_*m*_, *α*_*s*_, *β*_*m*_, *β*_*s*_ respectively. The values of these parameters are given in Table 2. Examples of receptive fields of LGN neurons in the model are plotted in Supp. Fig. 1B. The heterogeneity in the LGN receptive fields is depicted in Supp. Fig. 1C.

In the simulations of Sup. Fig. 9, the preferred orientations of LGN neurons are chosen randomly with a distribution

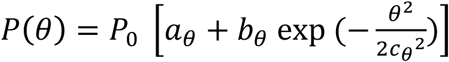

where *P*_0_ is a normalization constant. The parameters we used in these simulations are given in Table 3.

**Table 3:** Parameters of the computational model of mouse V1: elongated receptive fields of LGN cells.

|  |  |
| --- | --- |
| $\alpha_m$ | 0 |
| $\alpha_s$ | 0.2 |
| $a_\theta$ | 142 |
| $b_\theta$ | 1128 |
| $c_\theta$ | 21.5 deg |

(\*) Parameters of the log-normal distribution are normalized to obtain values in degrees.

## Supplementary Figures

**Supplementary figure 1:**
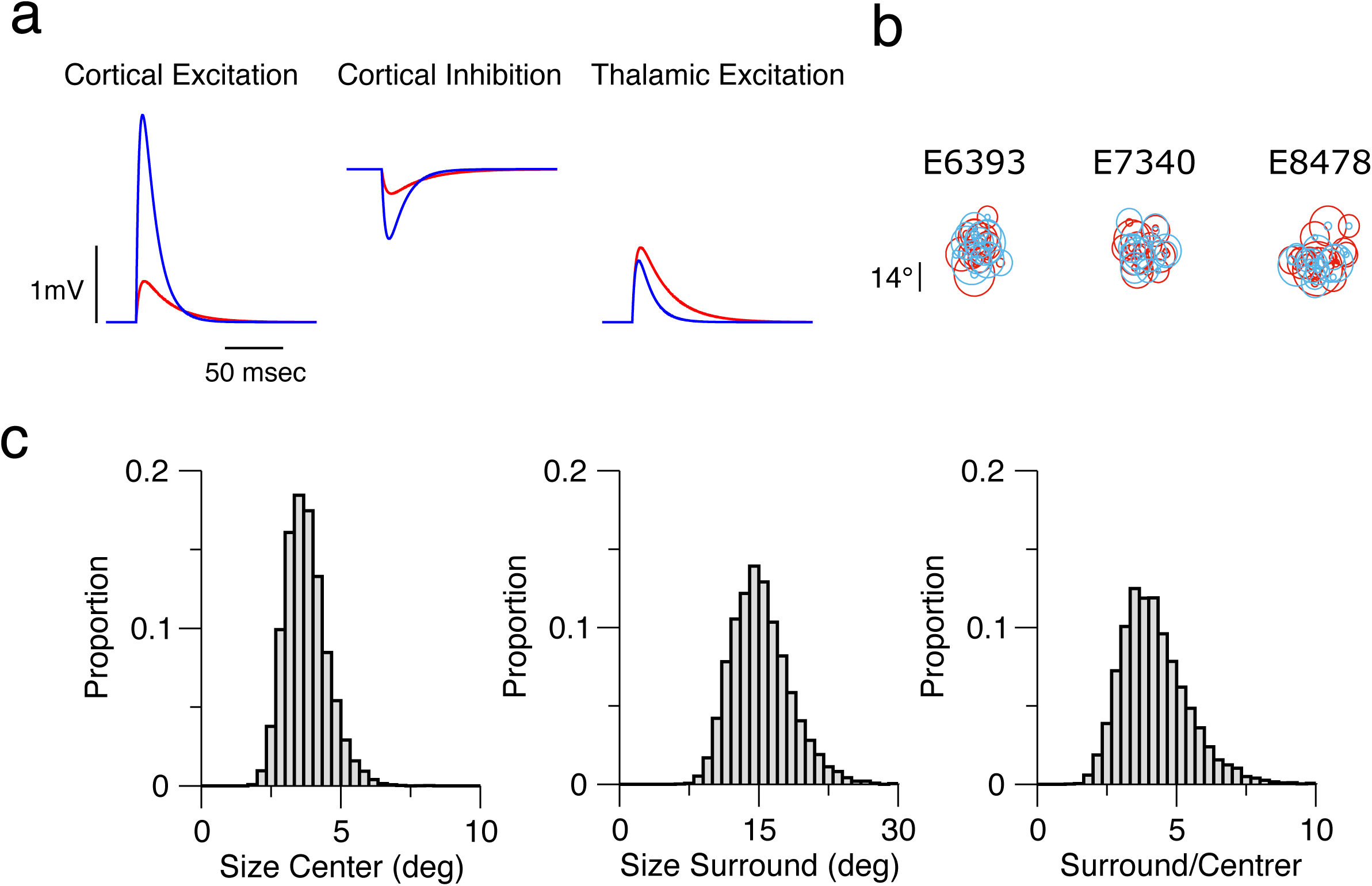
The large scale spiking model of V1. Parameters are as in Tables 1 and 2. **A.** PSPs for the cortical excitation (left), the cortical inhibition (middle) and the thalamic excitation (right). Red: PSPs for excitatory postsynaptic neuron. Blue: PSPs for inhibitory post-synaptic neuron. **B.** Receptive fields of thalamic neurons presynaptic to three representative cortical neurons. Thalamic neurons have circular receptive fields modeled as a difference of Gaussian describing OFF and ON subfields (see Methods). Red: ON subfield. Blue: OFF subfield. The radii of the plotted circles are the standard deviation (SD) of the corresponding Gaussians.. **C.** Histograms of the radii of center (left) and surround subfields (middle). Right: histogram for the ratio of the surround and the center subfields radii.

**Supplementary figure 2:**
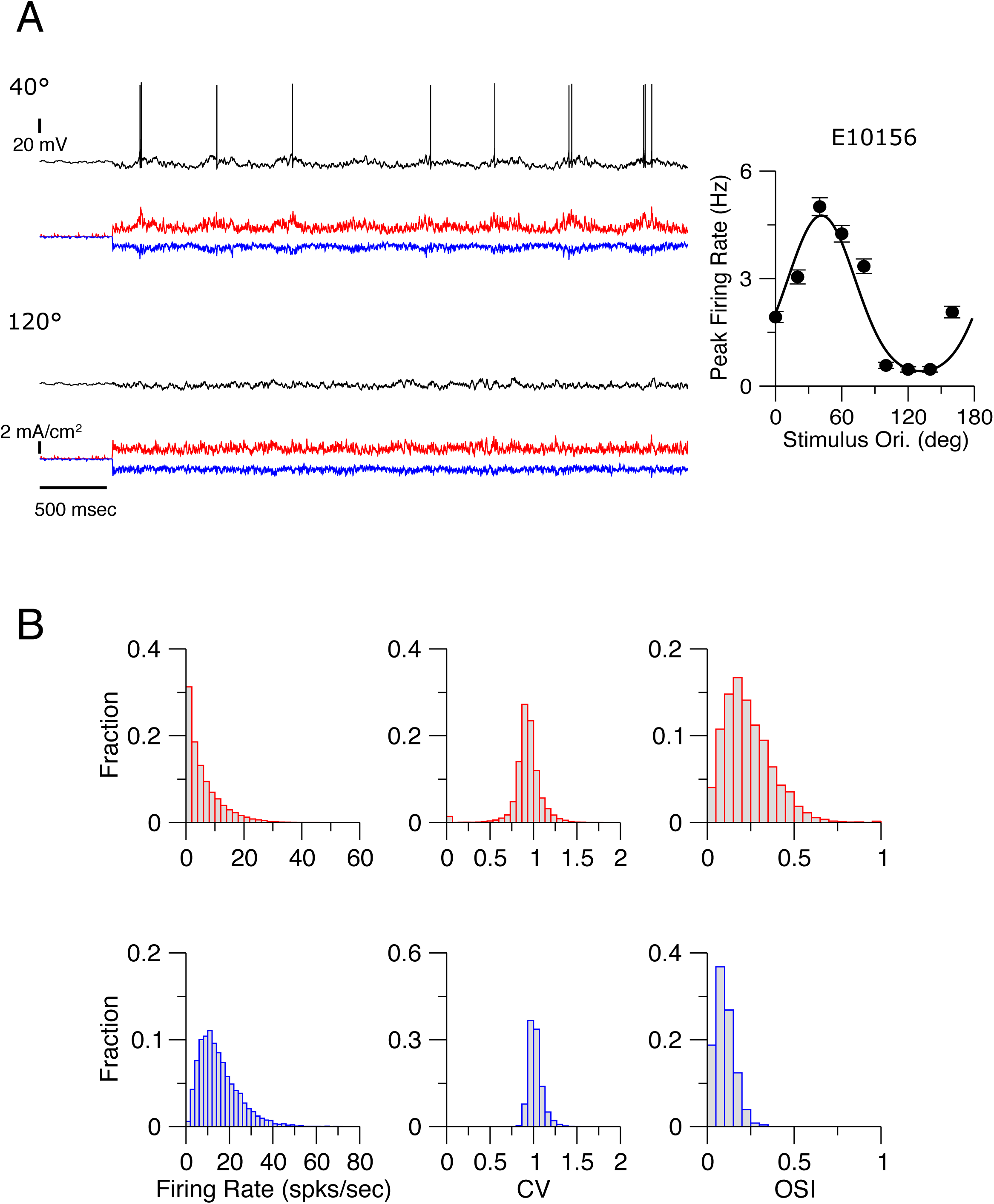
The V1 model network operates in the balances state. **A.** Voltage traces for one excitatory neuron. Stimulus begins at t=500msec. The drifting grating (SF = 0.03 cyc/deg) is presented at two orientations: 40° (upper trace) and 120° (lower trace). The panels below the voltage traces depict the excitation/inhibition balance. The excitatory (red), inhibitory (blue) currents to the neuron are plotted (note that in simulations all these currents and the voltage are always simultaneously known). Right panel: The tuning curve of the neuron. Average firing rate averaged over 80 sec. Parameters are as in Tables 1 and 2. **B.** Histograms of the peak firing rate (left), the coefficient of variation (CV) of the interspike distribution (middle). The stimulus is at 0° (SF=0.03 cyc/deg). Histograms are essentially the same for all stimulus orientations. The heterogeneity in firing rates and high temporal variability (CV around 1) of the neurons discharges are hallmarks of the balanced state. Right panel∷ The OSI of the peak response (SF=0.03 cyc/deg).. Red: Excitatory neurons. Blue: Inhibitory neurons. Inhibitory neurons are much less selective than excitatory neurons in agreement with experimental results (Kuhlman et al., 2011).

**Supplementary figure 3:**
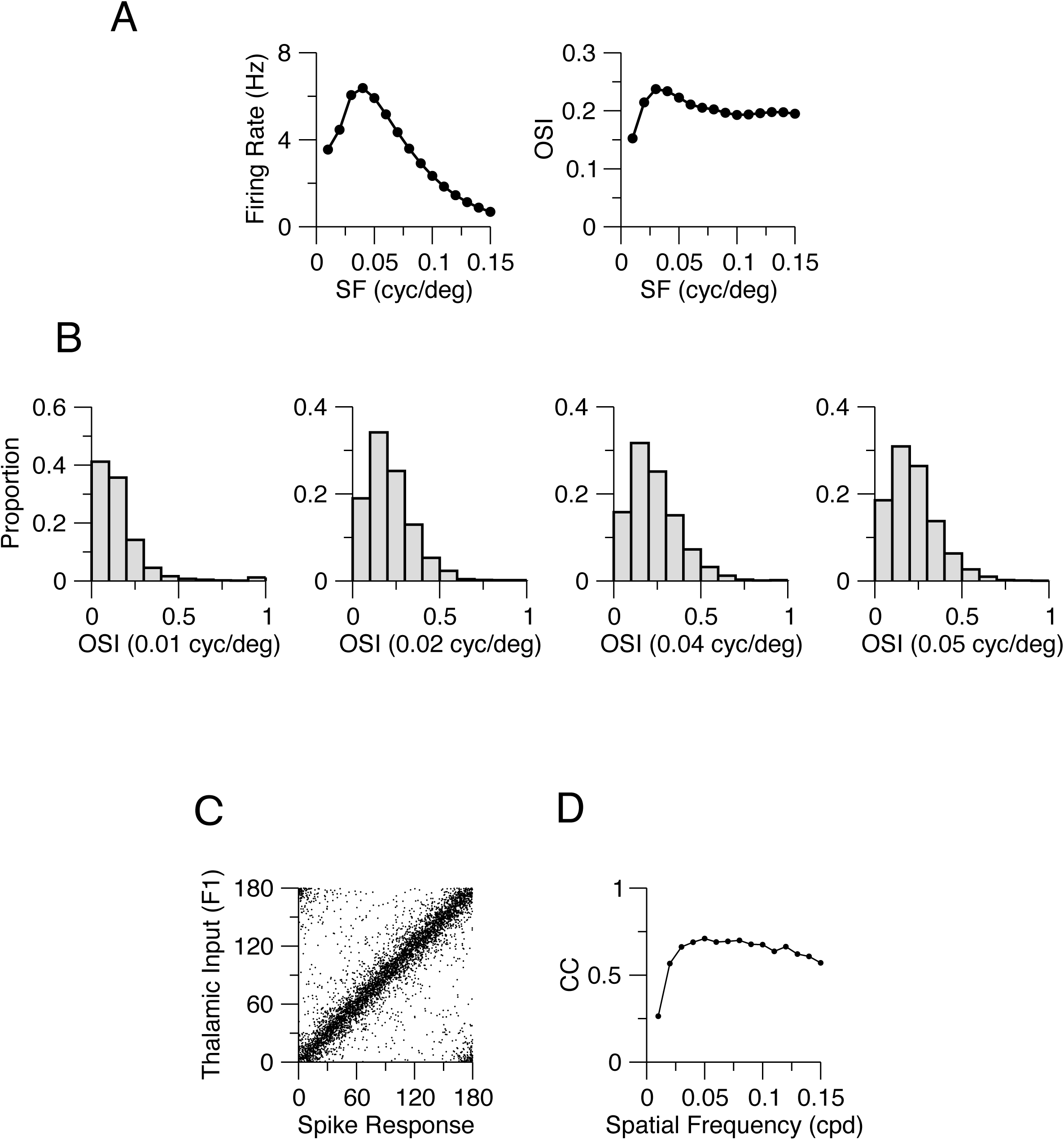
Dependence of the firing rate and selectivity on spatial frequency. **A.** Left: Population average of the peak firing rate at preferred orientation (E neurons) vs. grating spatial frequency (See Methods). Right: Population average OSI of the peak spike response vs. spatial frequency. B: Histograms of the OSI for different spatial frequency. (For 0.03 cycle/deg, see Fig. 3). The histograms are similar except for SF=0.01 cyc/deg. **C**: Preferred orientation of the thalamic excitation vs. preferred orientation of the spike response of the cortical neurons for SF=0.03 cyc/deg (n=5041). The correlation is strong. **D:** The circular correlation (CC, see Methods) of the thalamic input and spike response preferred orientations vs. the spatial frequency. The dependence on spatial frequency is weak except for low spatial frequency.

**Supplementary figure 4:**
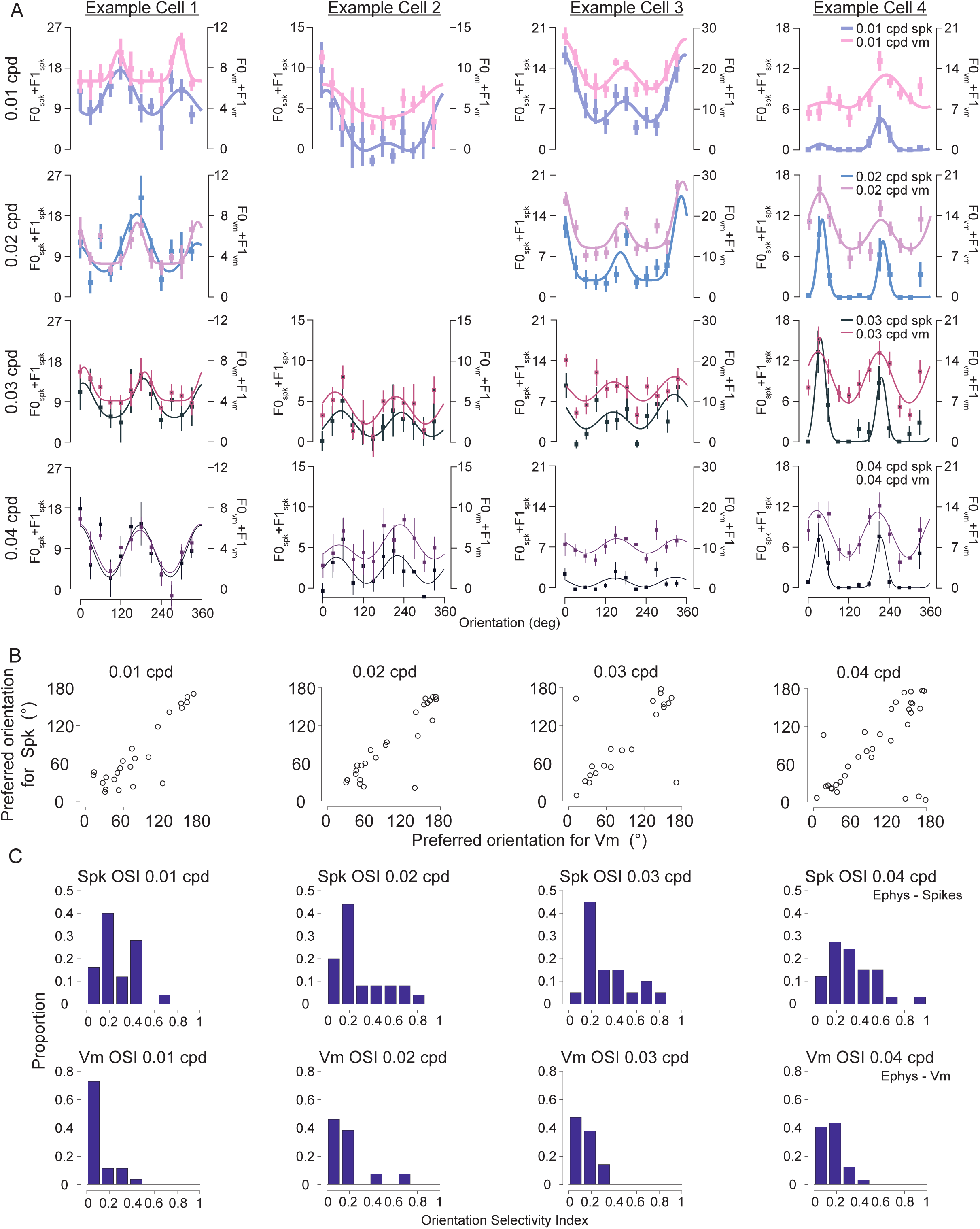
Vm and spike orientation preferences show similar dependency on spatial frequency. **A.** Each column represents a neuron. Each row is tuning curves for a different SF. Both the Vm and spikes based tuning curves show similar preference. **B.** The preferred orientation based on spikes vs. the preferred orientation based on Vm for all neurons. **C.** The orientation selectivity index (Methods) for spike rate (top row) and membrane potential (bottom row).

**Supplementary figure 5:**
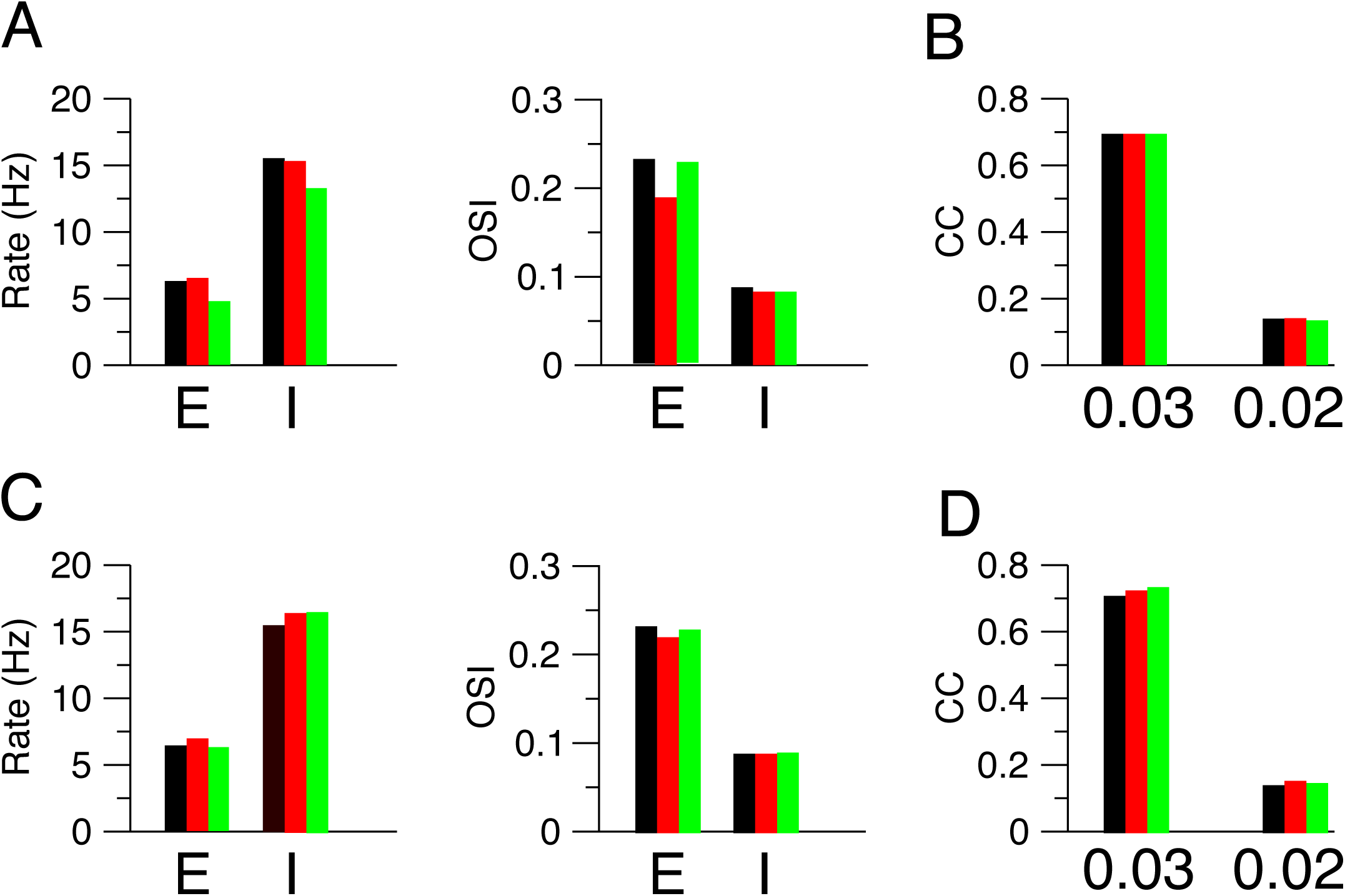
Robustness to changes in network size and connectivity parameters. **A**. Bar charts for the population averaged peak response (orientation of the stimulus: 0°; SF=0.04 cycle/deg) for excitatory (E) and inhibitory (I) neurons (left) and population averaged OSI (right). Black: Default set of parameters (Table 1,2). Red: N_E_=78400, N_I_=19600, N_L_=40000; other parameters as in Table 1, 2. Green: K_EL_=100; g_EL_ was increased to keep in both populations the firing rate approximately the same.: g_EL_=0.009 mS msec/cm^2^; other parameters as in Table 1, 2. **B.** Correlation coefficient (CC) for excitatory population between the preferred orientation of spike response for SF=0.04 cyc/deg and SF=0.03 cyc/deg (left) and SF=0.02 cyc/deg (right). For SF=0.01 cyc/deg: CC=0. Color code as in A. **C, D.** Same as in A,B but for changes in synaptic conductances. Black: Default values. Red: All conductances are multiplied by 0.75. Green: All conductances are multiplied by 1.25. Other parameters are as in Table 1,2.

**Supplementary figure 6:**
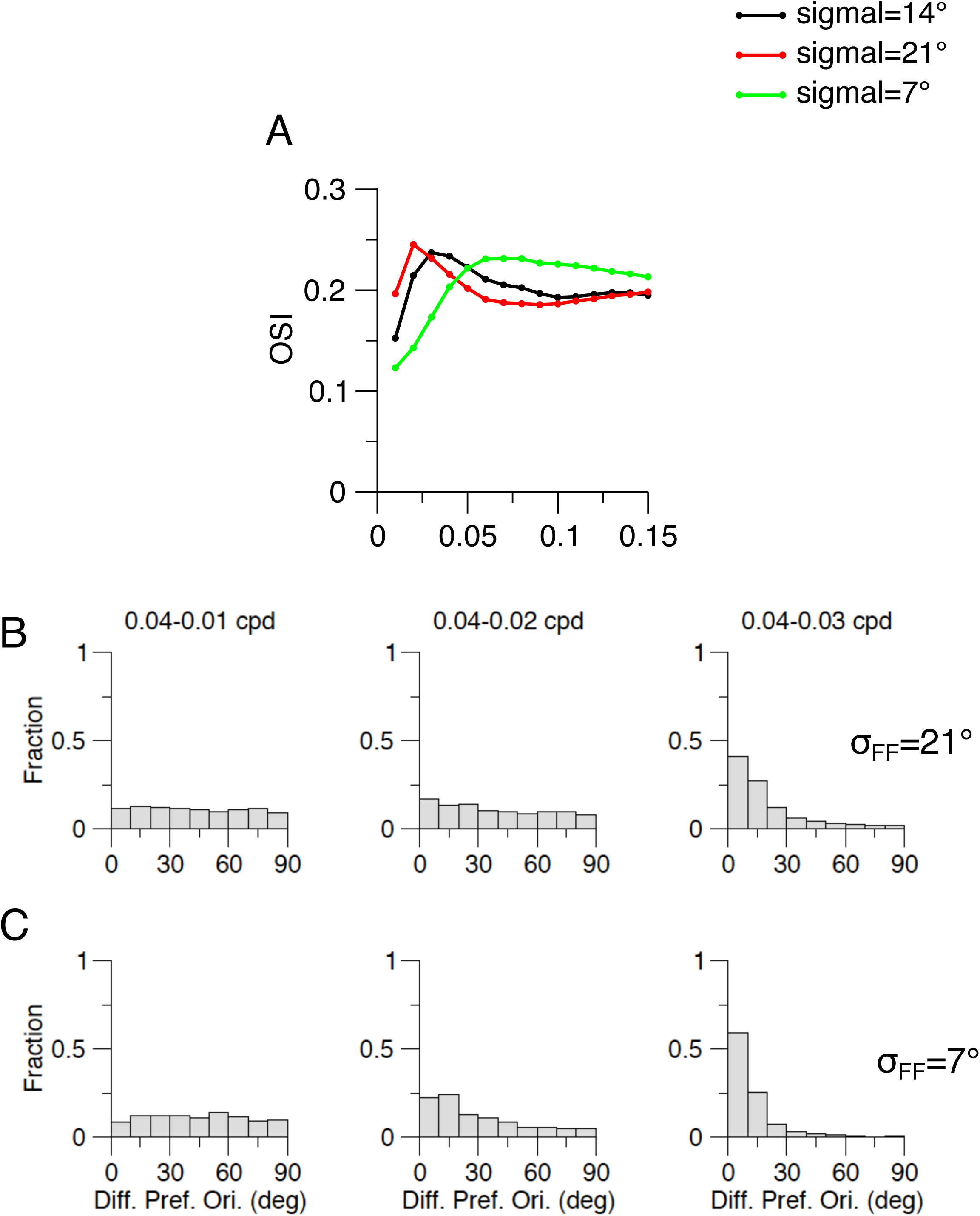
The decorrelation in orientation preference with spatial frequency depends on the dispersion of the thalamo-cortical projections. **A.** Population average OSI of the peak spike response of exciatory neurons vs. spatial frequency for σ_FF_. = 14° (black), 21° (red) and 7° (green). Other parameters as in Table 1, 2. **B.** Histograms of the difference in orientation preference between 0.04 cyc/deg and 0.01 (left), 0.02 (middle) and 0.03 (right) cyc/deg. Top: σ_FF_=7°. Bottom: σ_FF_ =21°. Compare with the corresponding histograms for σ_FF_=14° (Fig. 4). Only neurons with OSI larger than 0.2 for each pair of spatial frequencies are included. For σ_FF_=7°: CCs are 0.72 (0.04-0.03 cyc/deg), 0.18 (0.04-0.02 cyc/deg) and 0 (0.04-0.02 cyc/deg).. For σ_FF_=21°: CC=0.48, (0.04-0.03 cyc/deg), 0.03, (0.04-0.02 cyc/deg), 0 (0.04-0.01 cyc/deg).

**Supplementary figure 7:**
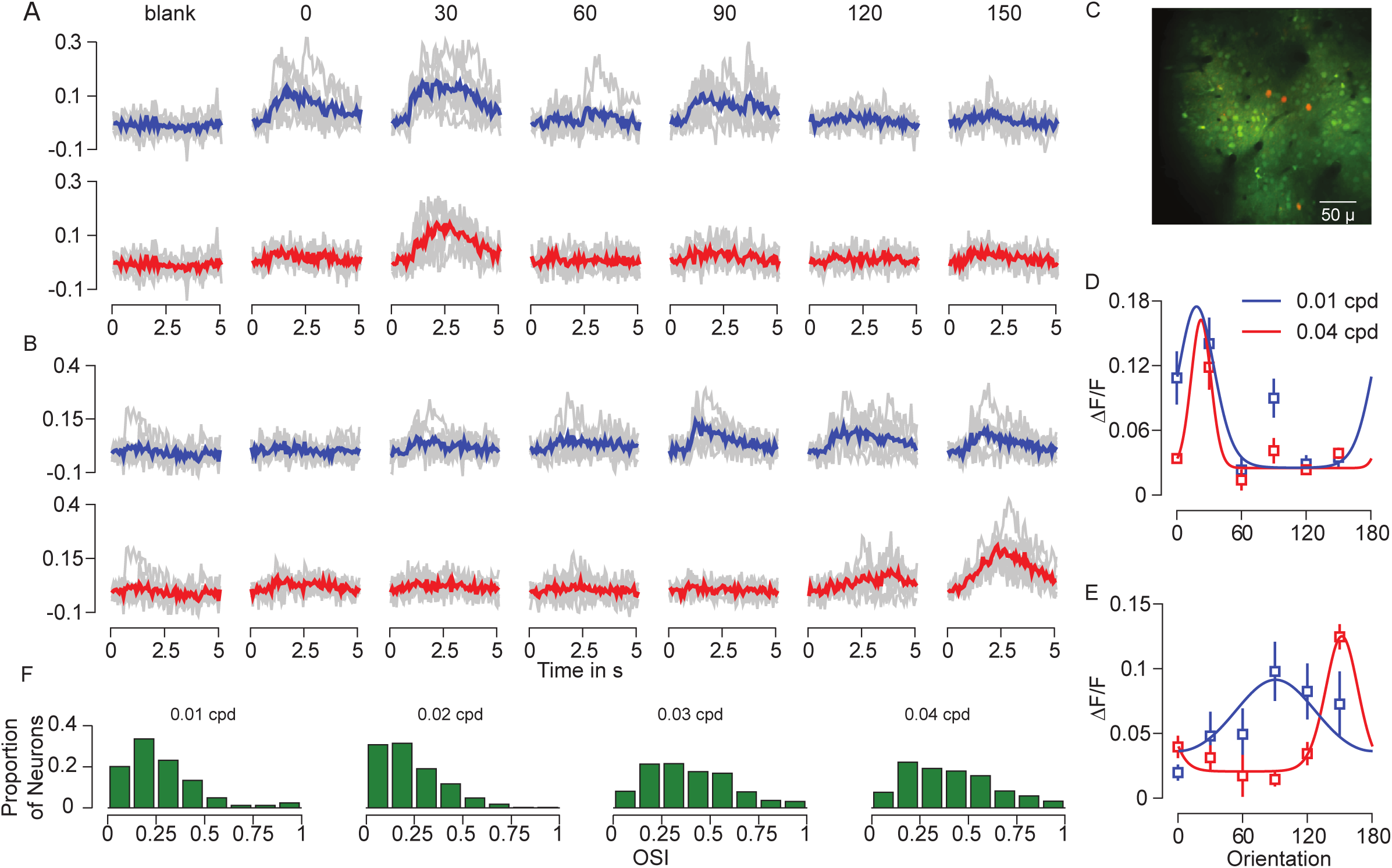
Example calcium orientation selectivity in mouse V1 A,B,.. The calcium responses for three example neurons for different orientation conditions and two spatial frequencies (0.01 cpd, blue and 0.04 cpd, red). **C.** Example imaging plane. **D, E**,. Orientation tuning curves for different SFs for the neurons represented in a and b. **F.** OSI distribution across all SFs used.

**Supplementary figure 8:**
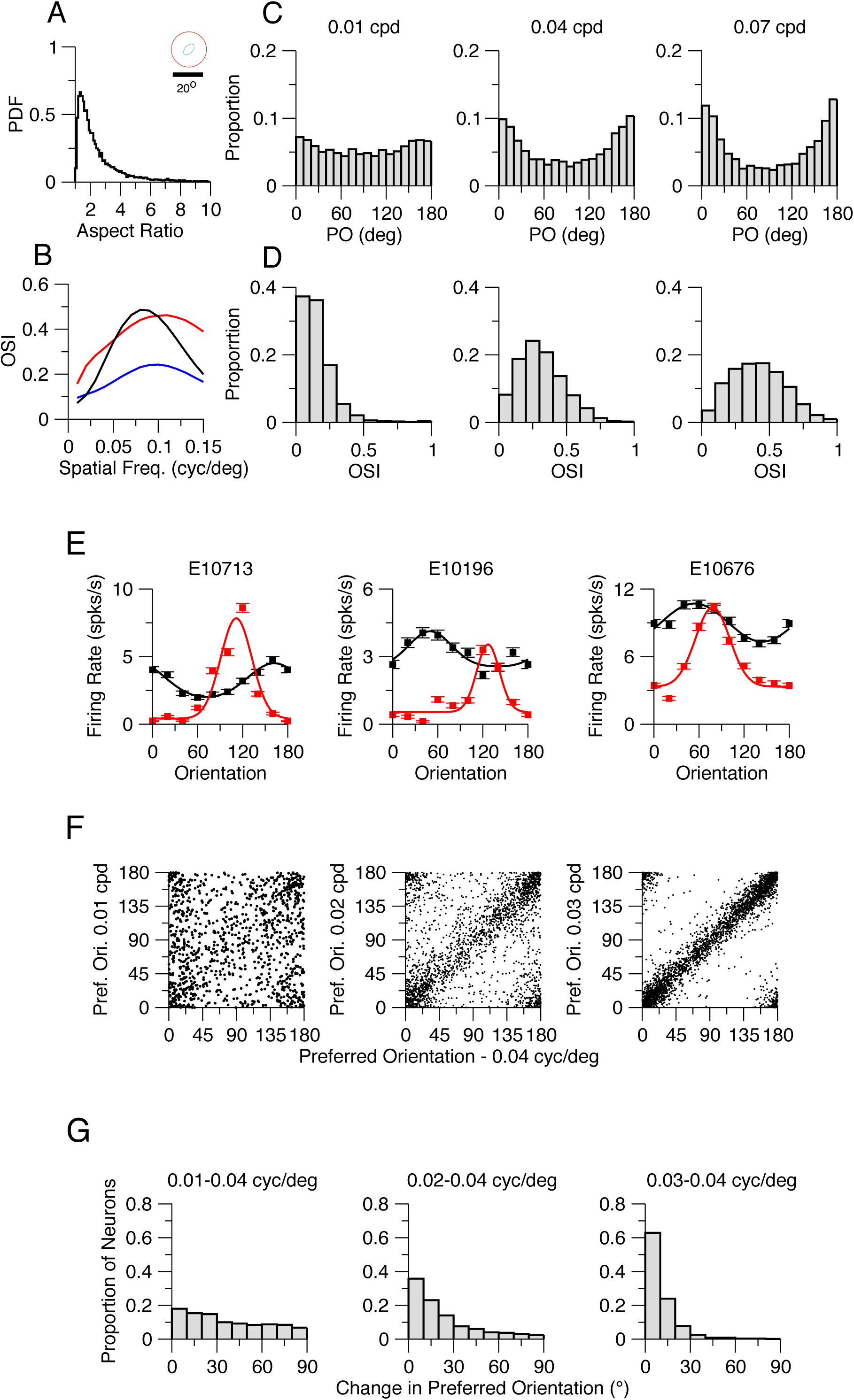
The V1 network model with elongated thalamic receptive. Parameters are as in Table 3. **A.** The central subfields of the receptive fields of LGN cells are elongated. Surround subfields are circular. Distribution of the aspect ratio for the center subfield is plotted.. Inset: Example of a thalamic neuron receptive field. **B.** Population average OSI for the spike peak response vs. spatial frequency. Red: E cells. Blue: I cells. Black: LGN cells. **C.** Histograms of the preferred orientations of the excitatory neurons for grating with 0.01 (left), 0.04 (middle) and 0.07 (right) cyc/deg. Preferred orientations are bias toward 0° because the distribution of the axis orientation of the center subfield of LGN cells is biased toward this orientation. **D.** Histograms of OSI for E neurons. Left to right: 0.01, 0.04, 0.07 cyc./deg. **E.** Tuning curves of three example E cortical neurons. Black: 0.01 cyc/deg; Red: 0.04 cyc/deg. **F, G.** Population data demonstrating the change in prefrerred orientation with spatial frequency. CC=0.77 (0.04-0.03 cyc/deg)., 0.33 (0.04-0.02 cyc/deg,) 0.04 (0.04 - 0.01 cyc/deg.

**Supplementary figure 9:**
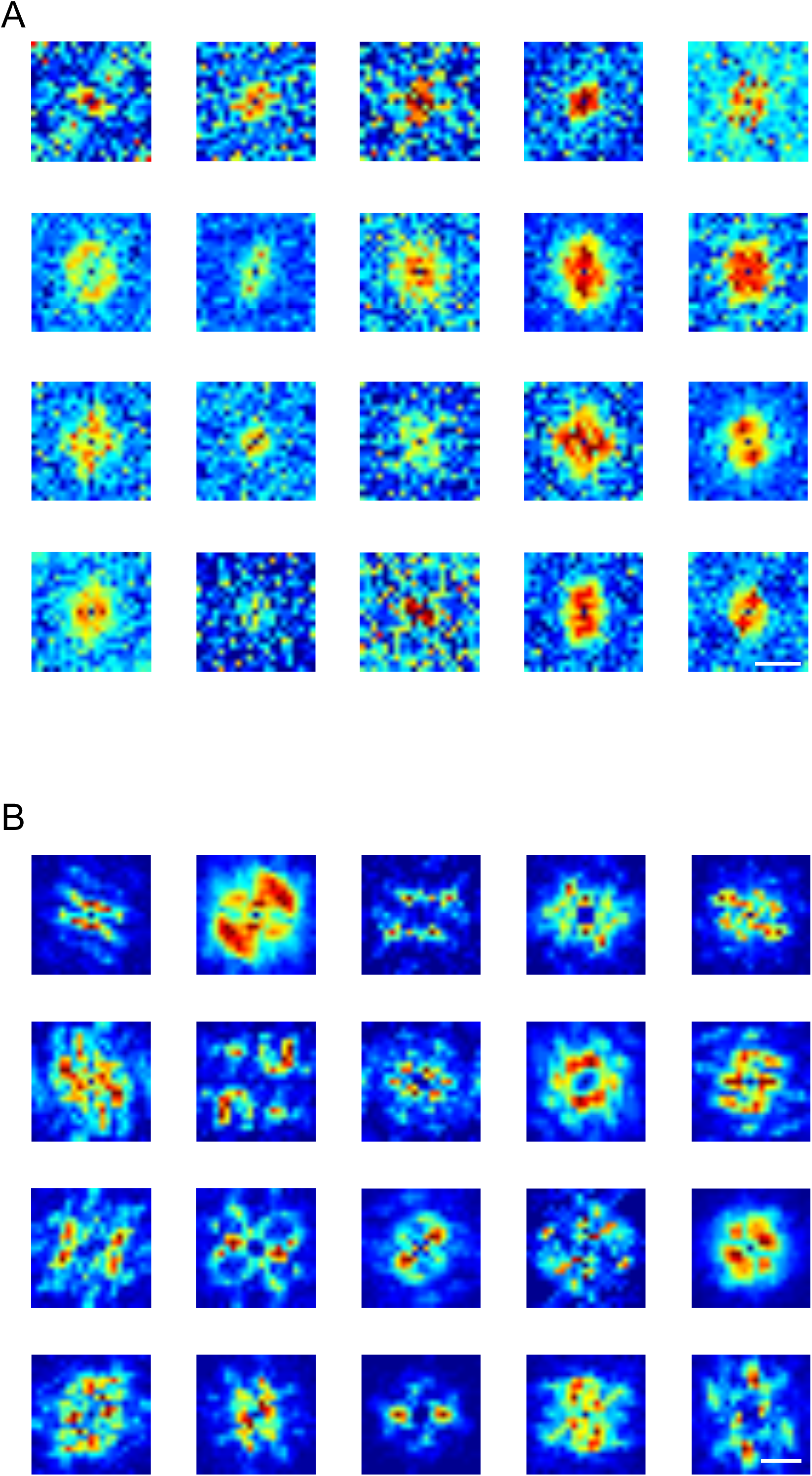
Examples of Hartley RFs based on electrophysiology and the model. **A.** 20 example neuron membrane potential responses to the Hartley stimulus. **B**. 20 example responses of model neurons to the Hartley stimulus.

**Supplementary figure 10:**
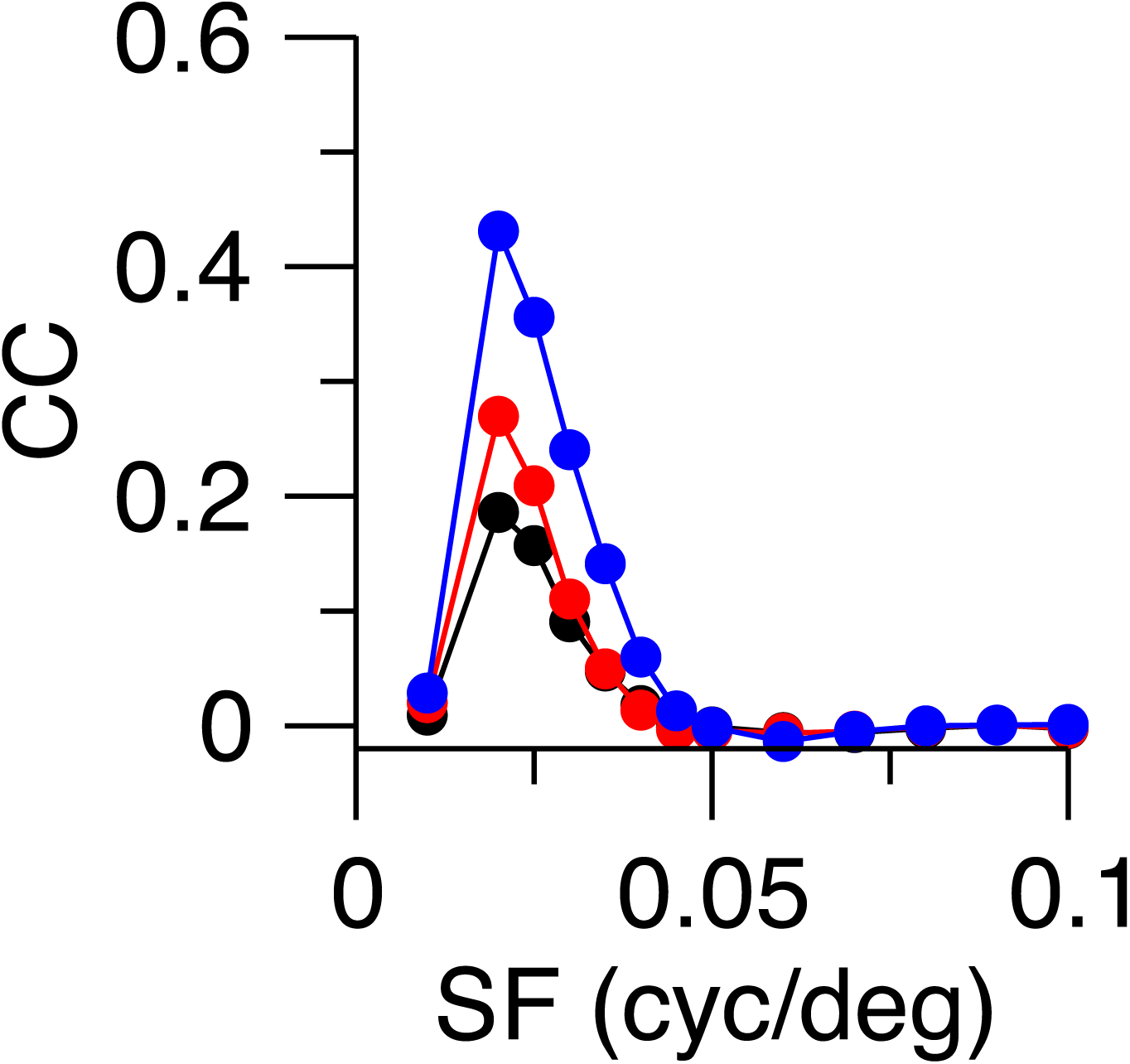
Correlation coefficient between the orientation of the ON-OFF offset axis and the orientation preference of the thalamic excitation vs. spatial frequency. Parameters of the model as in the default set. Black: All cells. Red: Offset > 2°. Blue: Offset > 4°.

